# A comprehensive approach to artifact-free sample preparation and the assessment of mitochondrial morphology in tissue and cultured cells

**DOI:** 10.1101/2021.06.27.450055

**Authors:** Antentor Hinton, Prasanna Katti, Trace A. Christensen, Margaret Mungai, Jianqiang Shao, Liang Zhang, Sergey Trushin, Ahmad Alghanem, Adam Jaspersen, Rachel E. Geroux, Kit Neikirk, Michelle Biete, Edgar Garza Lopez, Zer Vue, Heather K. Beasley, Andrea G. Marshall, Jessica Ponce, Christopher K. E. Bleck, Innes Hicsasmaz, Sandra A. Murray, Ranthony A.C. Edmonds, Andres Dajles, Young Do Koo, Serif Bacevac, Jeffrey L. Salisbury, Renata O. Pereira, Brian Glancy, Eugenia Trushina, E. Dale Abel

## Abstract

Mitochondrial dynamics (fission, fusion, and the formation of nanotunnels) and morphology are very sensitive to the cellular environment. Mitochondria may be adversely affected by oxidative stress, changes in calcium levels, and hypoxia. Investigating the precise relationship between organelle structure and function requires methods that can adequately preserve mitochondria while providing accurate, quantitative measurements of morphological attributes. Here, we demonstrate a practical approach for preserving and measuring fine structural changes using two-dimensional, high-resolution electron micrographs. This approach is further applicable for three-dimensional volume renderings, obtained using serial block-face and focused ion beam-scanning electron microscopy, highlighting the specific advantages of these techniques. Additionally, this study defines a set of quantifiable metrics that can be applied to measure mitochondrial architecture and other organellar structures. Finally, we validated specimen preparation methods that avoid the introduction of morphological artifacts that may interfere with mitochondrial appearance and do not require whole-animal perfusion.

## Introduction

Mitochondria are responsible for meeting the energetic and metabolic demands required for cellular changes^1^. The accurate estimation of mitochondrial size, shape, network organization, and interactions with other organelles could contribute to understanding disease mechanisms and the monitoring of therapeutic efficacy^2–12^. Mitochondrial number and shape are determined in part by fusion and fission cycles^2–7^. The fusion process is coordinated by the fusion proteins optic atrophy 1 (OPA1) and mitofusin-1 and -2 (MFN1 and MFN2, respectively)^8,13,14^. Meanwhile, the fission machinery includes dynamin-related protein 1 (DRP-1) and its receptors mitochondrial fission 1 protein (FIS1) and mitochondrial fission factor (MFF), and the mitochondrial dynamics proteins MiD49 and MiD51^5,7,8,10,14–16^.

Mitochondria interact with other intracellular organelles, including the endoplasmic reticulum (ER), through direct mitochondria–ER^1,17–23^ contacts (MERCs). Mitochondria also interact with lipid droplets (LDs), through either transient, kiss-and-run contacts or the longer-lived LD-anchored mitochondria (LDAM) contacts^2^. Mitochondria further communicate with the nucleus via the release of metabolic intermediates that mediate transcriptional regulation^1,4,17–24^. Furthermore, mitochondria form dynamic branching networks and connections via nanotunnels or the mitochondria-on-a-string (MOAS) morphology^25–28^. Nanotunnels and MOAS link mitochondrial elements together in response to energetic stress, hypoxia, and other physiological and environmental changes. Nanotunnels and MOAS are often detected during disease states, such as Alzheimer’s disease^25,29,30^. Their formation is thought to represent a mechanism for promoting mitochondrial communication and protecting mitochondria against fragmentation and lysosomal degradation^25–27,31,32^. These unique structures are often formed within minutes of exposure to hypoxic conditions. However, the methods employed during specimen preparation for electron microscopy (EM) could potentially expose cells and tissues to hypoxic conditions. Therefore, the development of methodologies capable of producing artifact-free specimens is imperative to allow for the reliable and reproducible assessment of changes in mitochondrial morphology. Given great complexity of mitochondrial morphological changes and intracellular interactions, the development of standardized approaches for quantifying these changes in a consistent manner will promote the reproducible evaluation of multiple samples for comparison across laboratories.

Since the first time mitochondria were imaged using EM in 1953^33,34^, myriad of methods have been developed for mitochondrial fixation, dehydration, sectioning, and staining, aiming to better preserve the native organellar morphology^24,35–42^. However, no clear consensus has been reached regarding which tissue or cell preparation methods are most likely to ensure reliable, reproducible, and artifact-free resolution of mitochondrial morphology using EM. Early studies of mitochondrial structure were limited to 2-dimensional (2D) optical or EM imaging techniques. More recently, specialized instrumentation has been employed to generate high-resolution volume renderings using 3-dimensional (3D) EM. The novel MOAS phenotype was only recently discovered using 3D EM reconstruction and is not obvious when using conventional transmission EM (TEM)^25^. Until this report, no systematic approach has been defined for the characterization of 3D mitochondrial morphology or the identification of MOAS in multiple tissue types. Methodical quantification and definition of the mitochondrial architecture remains necessary and may reveal mechanistic features that underlie various cellular events in clinical or experimental samples. Novel structures that have yet to be defined may be uncovered as the technologies used for EM reconstructions continue to evolve. Therefore, it is critical that a pragmatic approach to 3D microscopy techniques is developed and standardized.

The objectives of this study were to (1) develop an optimized approach for specimen handling and fixation that preserves mitochondrial morphology in cells and tissue for EM evaluation; (2) develop an optimized methodology for quantifying organellar characteristics using TEM micrographs and 3D EM reconstructions; and (3) validate the reproducibility of these methods for use in describing morphological changes that occur in subcellular organelles. We present comprehensive protocols that can be used to standardize the quantification of mitochondrial morphology. Specifically, we focused on quantifying mitochondrial morphological indices^2–7^, such as hyperbranching, mitochondrial volume, cristae morphology^8–12^, and the length and percentage coverage of MERCs^1,17,19,20,43^. Additionally, we optimized specimen preparation conditions to ensure the generation of artifact-free samples derived from cells and tissue. We applied 3D EM reconstruction, using manual serial section TEM, automated serial block-face (SBF)-scanning EM (SEM), or focused ion beam (FIB)-SEM, and aligned image stacks to observe 3D mitochondrial morphological changes. Tools including ImageJ (FIJI) and Reconstruct, widely available, open-source image analytical platforms, were used to quantify EM micrographs and display 3D images following segmentation, respectively^25,44,45^. In addition, proprietary software, including Ilastik^1^ and Amira^46^, were used for the reconstruction of FIB-SEM and SBF-SEM acquisitions, respectively. We examined the effects of gene deletion for known mitochondrial dynamics proteins, including OPA1, MFN2, and DRP-1, to validate our approach. This allowed us to evaluate which methodologies provide appropriate conditions for the reliable and reproducible monitoring of mitochondrial dynamics.

## Results

### Artifact-Free Preparation of Cultured Cells for Ultrastructural Studies Using EM

We first aimed to establish the optimal conditions for preserving mitochondrial ultrastructure of cultured cells during TEM examination, including the fine details of mitochondrial cristae morphology. The development of artifact-free preparation methods that allow for the evaluation of mitochondrial morphological changes under experimental conditions is vital. Insulin stimulation has been shown to increase mitochondrial fusion in cultured cardiomyocytes and skeletal muscle cells, resulting in larger, fused mitochondria with increased cristae density^5^. We stimulated myotubes prepared from differentiated skeletal muscle satellite cells with insulin for 2 hours prior to fixation. This was done to increase the mitochondrial size and cristae density and to provide a model for the validation of specimen preparation methods under these experimental conditions. We, then, compared eight different fixation methods featuring various cell harvesting and fixation techniques to determine the method that best preserved mitochondrial and cristae morphology (Figure 1A–H).

**Figure 1.**
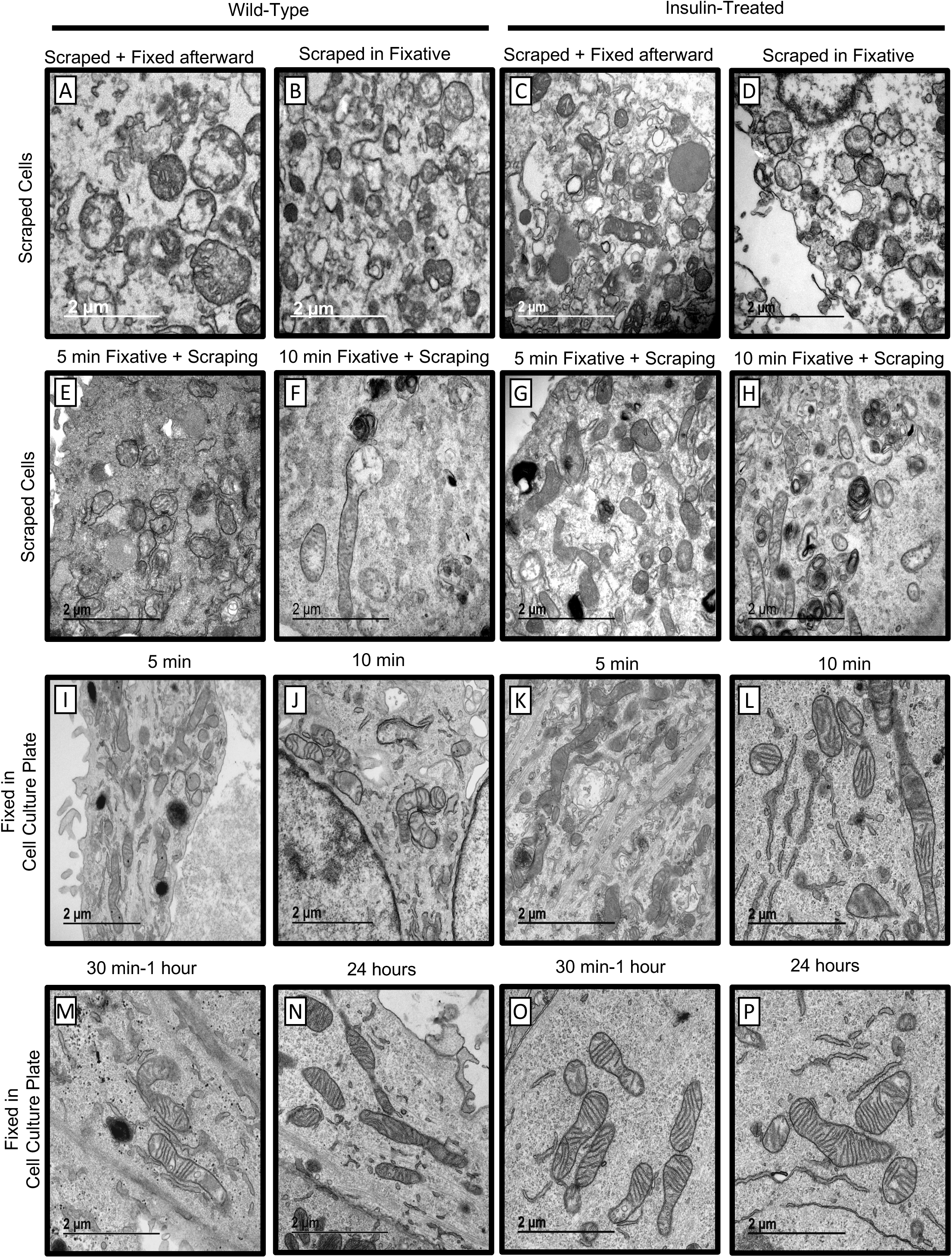
Comparison of the impact of various methodologies for fixing cultured cells on mitochondrial morphology determined by electron microscopy. Representative micrographs of cultured cells treated without and with insulin (**A-P**). **A, C.** Scraping followed by fixation. **B, D.** Scraping into transmission electron microscopy (TEM) fixative. **E, G.** Scraping after 5 minutes in TEM fixative. **F, H.** Scraping after 10 minutes in TEM fixative. **I, K.** 5 minutes in TEM fixative without scraping. **J, L.** 10 minutes in TEM fixative without scraping. **M, O.** 30–60 minutes in TEM fixative without scraping. **N, P.** 24 hours in TEM fixative without scraping. Insulin-treated cells were treated with insulin for 2 hours before fixation (**C-D, G-H, K-L, O-P**).

#### **•** Scraping with Fixation

Skeletal muscle cell cultures were grown in Matrigel-coated 6-well plates (Figures 1A–H). Following stimulation with or without insulin (2 hours at 10 nmol/L)^2^, treated myotubes were scraped with a standard cell scraper and post-fixed for TEM (Figure 1A, 1C). Scraping cells before fixation altered the plasma membrane and damaged mitochondria, resulting in the appearance of punctures in the membranes. Similarly, cells scraped immediately into TEM fixative (Figure 1B, 1D) or after fixation for 5 (Figure 1E, 1G) or 10 minutes (Figure 1F, 1H) were poorly preserved for TEM analysis. Therefore, we concluded that scraping during cell harvesting damages the plasma membrane and disrupts mitochondrial integrity. Therefore, we pursued alternative fixation and cell harvesting methods.

#### **•** Fixation without Scraping

Skeletal muscle cell cultures were grown in Matrigel-coated 6-well plates (Figures 1I–P). As opposed to the previous method, the TEM fixation and processing procedure was performed directly on the growth plate. A jigsaw was used to cut out the resin in the cell culture plate before completing the procedure. Once 80% confluency was achieved, the cultures were fixed by the addition of Trump’s fixative directly to the 6-well plates, followed by incubation at 37 °C for 5 (Figure 1I, 1K), 10 (Figure 1J, 1L), or 30–60 minutes (Figure 1M, 1O) or 24 hours (Figure 1N, 1P). The results showed that mitochondria were poorly resolved after 5 minutes of fixation (Figure 1I, 1K). Fixation for 10 minutes resulted in a slight improvement in mitochondrial morphology and cristae integrity (Figure 1J, 1L). However, fixation for 30–60 minutes (Figure 1M, 1O) greatly improved image quality, with no further improvements observed following fixation for 24 hours (Figure 1N, 1P). Low fixation times, both with and without insulin treatment, show poorly resolved cristae and fragmented mitochondria. Together these indicate that fixation without scraping for at least 1 hour was the most effective and efficient fixation method regardless of insulin treatment.

### Systematic Quantification of Mitochondrial Morphology and Interactions with Other Organelles

After determining the optimal fixation method, we sought to validate this method across multiple experimental conditions. The specific details regarding the methodology applied for the quantification of mitochondrial morphology in TEM images are presented in the online methods. The results obtained using these methods are presented to describe the observed changes in mitochondrial morphology and their interactions between themselves and other organelles in response to hormonal and genetic manipulations.

### Measuring Changes in Cristae Morphology Following OPA1 Ablation in Skeletal Muscle Myoblasts

OPA1 is a mitochondrial GTPase that maintains cristae morphology and is responsible for inner mitochondrial membrane fusion^14,47^. Mouse primary myoblasts in which *Opa1* levels were knocked down had decreased mitochondrial area (Figure 2A–C). Cristae morphology was also altered in these myoblasts, as demonstrated by reduced cristae score, cristae number, cristae volume, and cristae surface area compared with control cells expressing wildtype levels of *Opa1* (Figure 2D–G). Furthermore, compared with control cells, *OPA1*-deficient myoblasts demonstrated an increase in tubular cristae and a decrease in lamellar cristae (Figure 2H).

**Figure 2.**
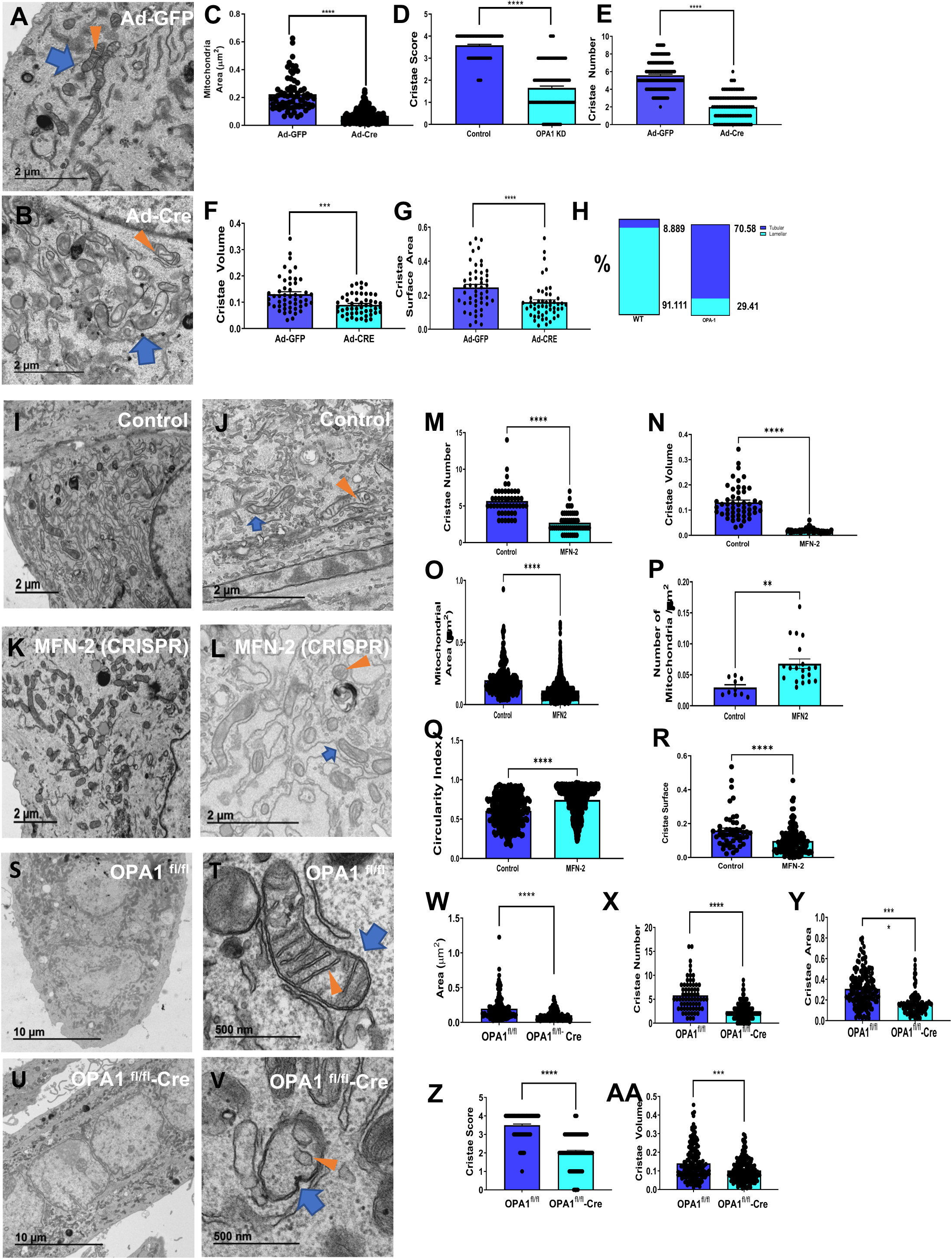
Quantification of mitochondrial characteristics in *OPA1*-deficient skeletal muscle myoblasts and *MFN2*-deficient or *OPA1*-deficient skeletal muscle myotubes. Blue arrows indicate mitochondria; orange arrows indicate cristae. Ultrastructure of (**A**), control and (**B**), *OPA1*-deficient primary skeletal muscle myoblasts. **C.** Mitochondrial area, **D.** Cristae score, **E.** Cristae number, **F.** Cristae volume, and **G.** Cristae surface area in control and *OPA1*-deficient primary skeletal muscle myoblasts. **H.** Percentage of tubular and lamellar cristae in control and OPA-1 deficient primary skeletal muscle myoblasts. Ultrastructure of **I-J,** control and **K-L,** *MFN2-*deficient primary skeletal muscle myotubes. **M.** Cristae number, **N.** Cristae volume, **O.** Mitochondrial area, **P.** Number of mitochondria per µm^2^, **Q.** Circularity index, and **R.** Cristae surface area in control and *MFN2*-deficient skeletal muscle myotubes. Ultrastructure of **S-T** control and **U-V** *OPA1*-deficient primary skeletal muscle myotubes**. W.** Area, **X.** Cristae number, **Y.** Cristae area, **Z.** Cristae score, and **AA.** Cristae volume in control and *OPA1*-deficient skeletal muscle myotubes. Scale bars: **A and B and I–L,** 2 µm; **S,U**, 10 µm; **T,V**, 500nm. Recombinant eGFP adenovirus (Ad-GFP) serves as a control while adenovirus-cre represents the experimental knockout. OPA-1 fl/fl represents a control while fl/fl-cre represents deletion of floxed OPA-1.

### Measuring Changes in Cristae Morphology Following *MFN2*, *OPA1, and DRP-1* Ablation in Skeletal Muscle Myotubes

MFN2 is a mitochondrial outer membrane GTPase responsible for mitochondrial outer membrane fusion^17,20^. Past research has shown cristae remodeling upon loss of MFN2^20,48,49^. We sought to verify this with our method and see if changes caused by MFN2 loss resemble changes caused by OPA1 loss. DRP-1 is a fission protein, which can be activated upon cellular stress, although its precise mechanism of action remains to be fully elucidated^16^. It also has important implications in calcium uptake, and prior research has shown that the loss of DRP-1 results in poor mitochondrial function and enlarged mitochondria^7^. We transiently ablated *Mfn2, OPA1, and DRP-1* in primary myotubes using the clustered regularly interspaced short palindromic repeat (CRISPR)-CRISPR-associated (Cas)9 system and used western blotting and quantitative PCR to verify their successful ablation (SFigure 1A-D). Compared with wild-type (WT) control myotubes (Figure 2I–L), *Mfn2*-ablated primary myotubes were characterized by reductions in the cristae number, cristae volume, mitochondrial area, and cristae surface area (Figure 2M–O, 2R). In contrast, the number of mitochondria and the circularity index increased (Figure 2P–Q). The results observed following ablation of *Opa1* from primary myotubes (differentiated muscle cells) using Cre recombinase (Figure 2S–V) were similar to results observed in *Opa1* knockdown myoblasts (undifferentiated muscle cells). Loss of *Opa1* in myotubes resulted in a decrease in the mitochondrial area, cristae number, cristae area, cristae score, and cristae volume (Fig. 2W–AA). In quantifying the mitochondria upon loss of *DRP-1* (SFigure 1E-J), we observed an increase in mitochondria length and mitochondria area (SFigure 1K-L). For cristae, while the cristae score and surface area decreased, signifying less well formed and smaller cristae, the number of cristae increased (SFigure 1M-O). Importantly, with optimal fixation as described above, we further elucidated^20,50^ a potential role for OPA1 in MERC architecture. We also sought to further elucidate the specific effect of MFN2 and DRP-1 on MERC architecture.

### Determining MERC Distance in Murine *MFN2*-Deficient Cells, *OPA-1*, *DRP-1* Ablated Myotubes

Although several studies have reported that the ablation of *Mfn2* alters MERC distance^6,11,12^, other studies have disputed this conclusion^17,51^. Because these discrepancies may be due to suboptimal fixation methods^13,15,23,51–53^, we sought to measure the distances between rough ER and mitochondria in *MFN2*-deficient cells using our optimized fixation protocol. Furthermore, we also examined DRP-1, which similar to MFN2 may could be subjected to post-translational modification on MERC tethers^54^. Not all ER-mitochondria contacts are direct, therefore, it is important to use multiple quantification approaches when measuring MERCs to gain a complete picture. This includes distance between ER and mitochondria, and percent coverage between the two. In quantifying MERCs, we used the definition of MERCs proposed by Giacomello & Pellegrini (2016) which classifies MERC width as typically around 10 to 50 nm^11^. Ablation of *MFN2* from fibroblast cells using Cre recombinase (Figure 3A–B) increased the MERC distance compared with WT fibroblasts (Figure 3C). Next, we looked at percentage coverage of MERCS at a particular surface. Percentage coverage measures the percentage of the total endoplasmic reticulum or the mitochondrial surface that represent MERCs. The percentage coverage of the mitochondrial surface area with ER and the percentage coverage of the ER surface area with mitochondria both increased in *MFN2*-knockout (KO) cells relative to control cells (Figure 3D– E)^6^. Ablation of *MFN2* from primary myotubes using the CRISPR-Cas9 system resulted in a similar increase in MERC distance compared with control cells (Figure 3F–H). We also performed ultrastructural quantifications in *OPA1*-deficient primary skeletal muscle myotubes (Figure 3I–J), which revealed that, contrary to what we observed following *Mfn2* deletion, *Opa1* deletion decreased MERC distance (Figure 3K). Furthermore, it increased the percent coverage of both the ER and mitochondria (Figure 3L–M) relative to control myotubes. We further performed quantification in DRP-1-deficient myotubes (SFigure 2A-D), which showed that, similar to Mfn2 deletion, there was an increase in MERC distance and area of distance between MERCs (SFigure 2E-F). These data suggest that MERC distance may not always correlate with coverage, which could be an index of ER stress, or be related to altered calcium transfer between ER and mitochondria^14^. These results verified that our optimized fixation method retained organellar and sub-organellar features following gene knockdown.

**Figure 3.**
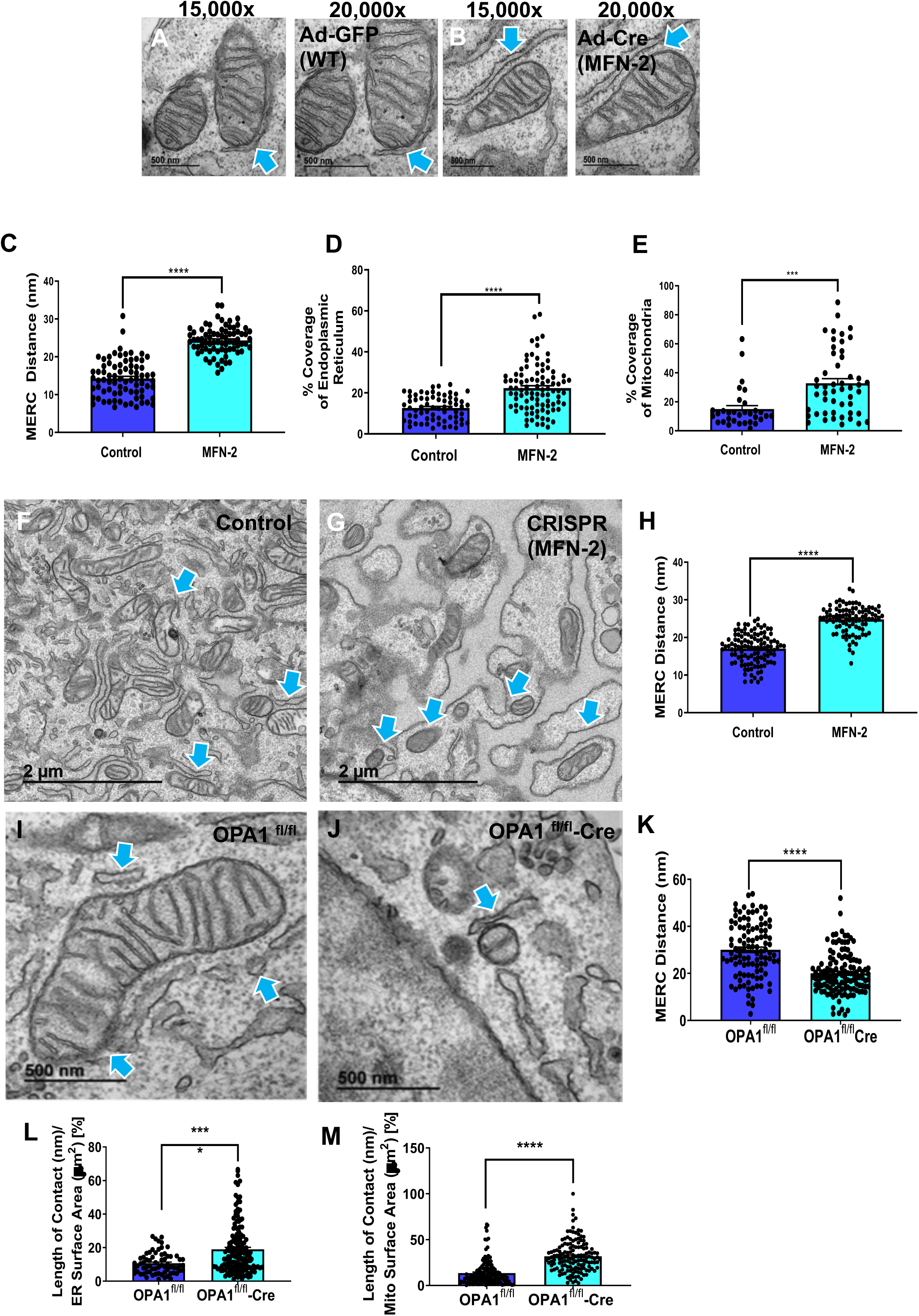
Mitochondria-Endoplasmic Reticulum Contact Site (MERC) ultrastructure in *MFN2*-deficient primary fibroblasts and *OPA1*-deficient primary skeletal muscle myotubes. **A and B.** Control and *MFN2*-deficient primary fibroblasts were imaged using transmission electron microscopy (TEM) at 15,000× and 20,000× magnification, then mitochondria–endoplasmic reticulum (ER) contacts (MERC) features were quantified. Light blue arrows indicate MERCs. **C.** MERC distance in control and *MFN2*-deficient primary fibroblasts. **D.** Percent ER coverage in control and *MFN2*-deficient primary fibroblasts. **E.** Percent mitochondria coverage in control and *MFN2*-deficient primary fibroblasts. Ultrastructure of **F,** control clustered regularly interspaced short palindromic repeats (CRISPR) and **G,** CRISPR-generated *MFN2*-deficient primary myotubes. Light blue arrows indicate MERCs. **H.** MERC distance in control CRISPR and *MFN2*-deficient primary myotubes. Ultrastructure of **I,** control and **J,** *OPA1*-deficient primary skeletal muscle myotubes. **K.** MERC distance, **L.** Length of contact (nm) per mitochondria surface area (µm^2^) percentage, and **M.** Length of contact (nm) per ER surface area (µm^2^) percentage in control and *OPA1*-deficient primary skeletal muscle myotubes. Scale bars: **A and B**, 500 nm; **F and G**, 2 µm; **I and J**, 500 nm. Recombinant eGFP adenovirus (Ad-GFP) serves as a control while adenovirus-cre represents the experimental knockout. OPA-1 fl/fl represents a control while fl/fl-cre represents deletion of floxed OPA-1.

### Artifact-Free Tissue Preparation for Ultrastructural Studies Using EM

To broaden the scope beyond cells, we sought to determine optimal techniques for fixation of tissue samples obtained from laboratory animals, in particular mice Commonly used techniques for collecting and fixing tissues for EM analysis include multiple methods of anesthesia, followed by cardiac perfusion. During cardiac perfusion, phosphate-buffered saline (PBS) buffer is flushed through the tissue to remove erythrocytes from blood vessels before fixative perfusion. Common anesthetics include ketamine/xylazine injections, CO_2_ inhalation, or inhalation of a 5% isoflurane/oxygen mixture. Cardiac perfusion ensures the thorough fixation of the whole body for the analysis of multiple tissues. However, this fixation method can prohibit the performance of other assays, such as western blot or RNA sequencing, which require fresh tissue. By contrast, other methods of euthanasia, such as CO_2_ or isoflurane inhalation followed by subsequent cervical dislocation, allow for the collection of multiple tissues for various analyses from the same animal, making these approaches attractive. To determine whether the use of any anesthetic agents induces hypoxic conditions, which could alter mitochondrial ultrastructure, we first compared mitochondrial morphology in hippocampal brain tissues obtained from wild type (WT) mice anesthetized with either CO_2_ (3 minutes) or 5% isoflurane/oxygen prior to cervical dislocation (Figure 4A–C). The whole brain was quickly removed, and hippocampal tissue (3 × 3 × 1 mm) was dissected and immediately immersed in Trump’s solution for fixation (Figure 4A). Consecutive serial tissue sections from animals subjected to each anesthetic technique were examined using TEM. We found that CO_2_ inhalation induced a prominent MOAS phenotype (Figure 4B). Tissue prepared following the inhalation of 5% isoflurane/oxygen did not exhibit significant MOAS formation, containing uniformly elongated mitochondria (Figure 4C).

**Figure 4.**
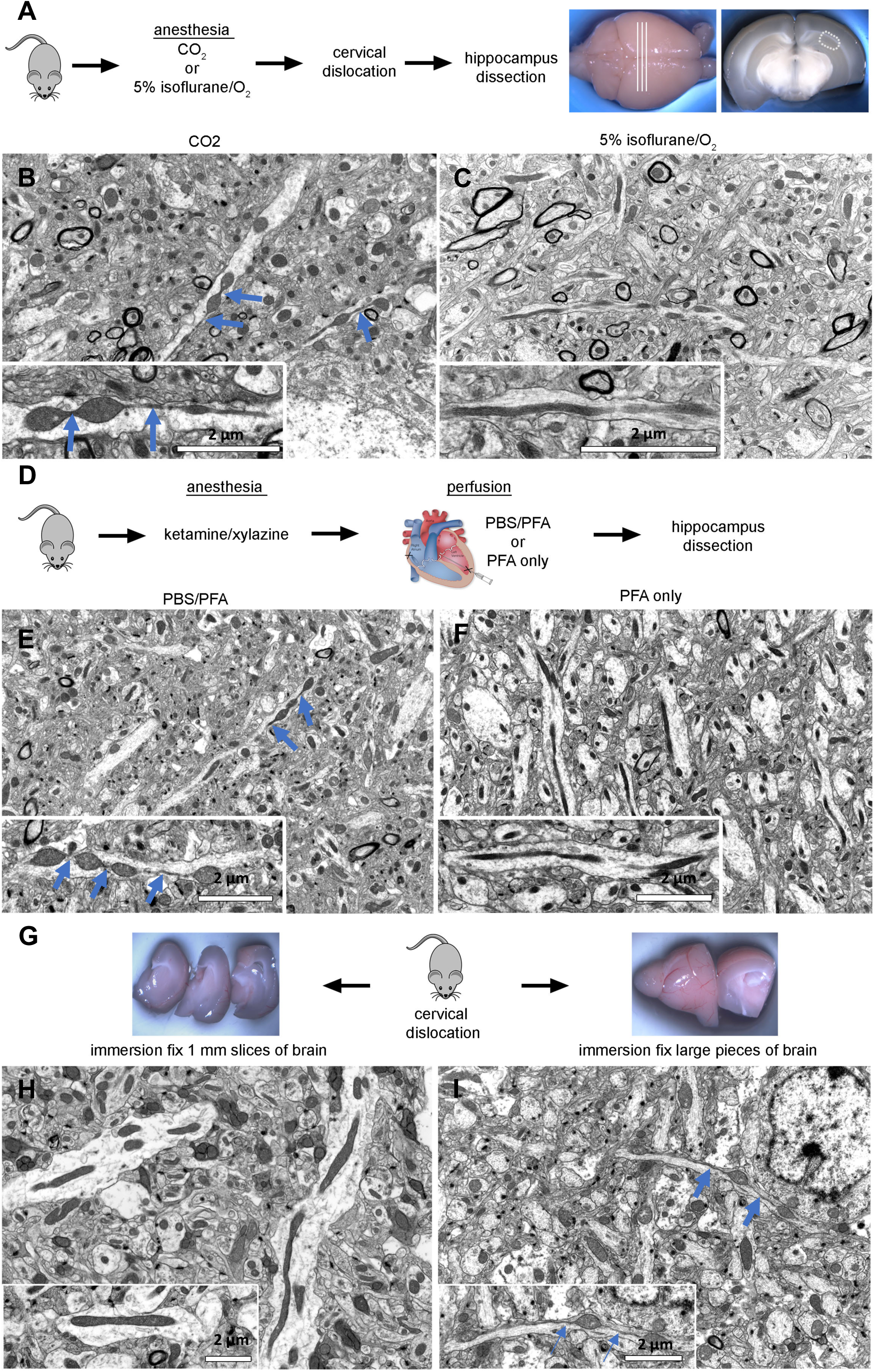
Evaluation of conditions allowing artifact-free tissue preparation to assay mitochondrial morphology using EM. **A.** Wild-type (WT) mice were sacrificed by cervical dislocation after anesthesia with CO_2_ or 5% isoflurane/oxygen inhalation. Fresh brains were removed and cut into coronal sections. The CA1 hippocampal region (hipp) from each section was dissected and further processed for transmission electron microscopy (TEM) or scanning electron microscopy (SEM). **B**. CO_2_ exposure for 3 minutes was sufficient to induce mitochondria-on-a-string (MOAS) formation in hippocampus tissue of WT mice. Scale bar, 2 µm. **C.** Mitochondria in hippocampus from mice administered 5% isoflurane prior to cervical dislocation and subsequent dissection maintained a normal shape and size typical for WT mice. Scale bar, 2 µm. **D.** WT mice were anesthetized using ketamine/xylazine, following by cardiac perfusion with either phosphate-buffered saline (PBS) flush and 4% paraformaldehyde (PFA) or 4% PFA only. Mice were euthanized by cervical dislocation, and the brains were removed and fixed in Trump’s solution overnight. The next day, brains were cut into coronal sections. The CA1 hippocampal region (hipp) from each section was dissected as shown in (**A**) and subjected to further processing for TEM or SEM. **E and F.** Cardiac perfusion preceded by PBS flush induced MOAS formation in hippocampal tissue of WT mice. Perfusion without PBS flush did not cause MOAS formation. Scale bar, 2 µm. **G.** WT mice were euthanized by cervical dislocation without prior anesthesia. The brains were removed, and one hemisphere was cut into 1-mm-thick slices (thin, left), while the other hemisphere was cut into two halves (thick, right). Tissues were subjected to immersion fixation in Trump’s solution overnight. Tissues cut in thin slices were free of MOAS (**H**), while larger pieces of tissue displayed pronounced MOAS formation **(I**). Scale bars, 2 µm. Blue arrows show nanotunnels.

We next examined how different methods of transcardial perfusion affect mitochondrial morphology in brain tissue (Figure 4D–F). WT mice were anesthetized with ketamine/xylazine injection and perfused with either PBS followed by 4% paraformaldehyde (PFA) or 4% PFA alone, without a PBS flush (Figure 4D). Although the PBS/PFA procedure typically requires approximately 5 minutes per mouse, perfusion with PFA alone can be performed in less than 3 minutes. Intact brains were removed and post-fixed overnight at room temperature in Trump’s solution. The hippocampal CA1 region and cortex were dissected from each brain and processed for TEM. The perfusion of animals with PBS before PFA fixation resulted in MOAS formation in hippocampus and cortex (Figure 4E, hippocampus is shown). By contrast, animals perfused with PFA alone without a prior PBS flush presented with uniformly elongated mitochondria throughout the examined brain regions (Figure 4F). These data suggest that transcardial perfusion with a PBS flush before the fixation may create hypoxic conditions that affect mitochondrial morphology.

The simple tissue immersion fixation technique is often used instead of a whole-body fixation by cardiac perfusion. The success of immersion fixation depends on the speed at which the fixative penetrates the tissue. If the tissue is too large, a slow fixation process could reduce tissue oxygenation, leading to hypoxia, which could alter mitochondrial morphology. To determine the optimum conditions for immersion fixation, we compared two sets of brain tissue obtained from the same WT mouse (Figure 4G). The mouse was euthanized by cervical dislocation without prior anesthesia, and the intact brain was rapidly removed (<2 minutes). One hemisphere was sliced into 1-mm-thick sections (Figure 4H), whereas the other was cut in half (Figure 4I). All tissues were immediately immersed in Trump’s fixative solution overnight. The hippocampal CA1 region and cortex were dissected from the 1-mm-thick tissue slices and from the middle of each half of the other hemisphere (Figure 4H-I) and processed for TEM. Mitochondria from the 1-mm-thick slices were approximately 0.3 µm in diameter and were uniformly elongated in all brain regions examined (Figure 4H, hippocampus is shown). By contrast, mitochondria from the larger brain tissue samples exhibited a wide variety of shapes, including MOAS, in all regions examined (Figure 4I, hippocampus is shown). These data suggest that the immersion fixation of large pieces of tissue may lead to altered mitochondrial morphology.

Taken together, our findings suggest that for accurate and artifact-free preservation of mitochondrial morphology in tissues prepared for TEM, suitable anesthetic agents include ketamine/xylazine or 5% isoflurane/oxygen. We found that CO_2_ should be avoided and if cardiac perfusion must be performed, a PBS flush should also be avoided. For the immersion fixation of fresh tissue, the tissue should not exceed 1 mm in thickness.

### 3D-EM Methodology

2D measurements were able to demonstrate alterations in the cristate morphology following *Opa1* and *Mfn2* loss. However, there are important mitochondrial architectural features that are not visible in 2D images but can be revealed using 3D EM techniques ^1,11,21–23,25,33,43,55^. Thus FIB- and SBF-SEM was used to further characterize mitochondrial ultrastructural and morphological changes following gene deletion. These advances in volume EM techniques, coupled with optimal preparation methods, have facilitated the 3D visualization of specimens with unparalleled detail, leading to increased interest in the ultrastructural investigation of cells and tissues. In general, serial section procedures, previously the domain of specialists, are becoming increasingly automatic through technologies including automatic tape-collecting ultramicrotomes and SBF- and FIB-SEM.

We tested whether the techniques developed for the preservation of mitochondrial morphology in brain tissue using 2D EM are suitable for 3D SBF-SEM. Serial-section TEM images (Figure 5A) from brain tissue of a mouse model of Alzheimer’s Disease, the APP/PS1 mouse^25^, were aligned in an image stack, and the Reconstruct software package was used to visualize the 3D mitochondrial morphology (Figure 5B). As demonstrated previously^25^, MOAS could clearly be identified in the neuropil of the hippocampus of the APP/PS1 mouse (Figure 5B).

**Figure 5.**
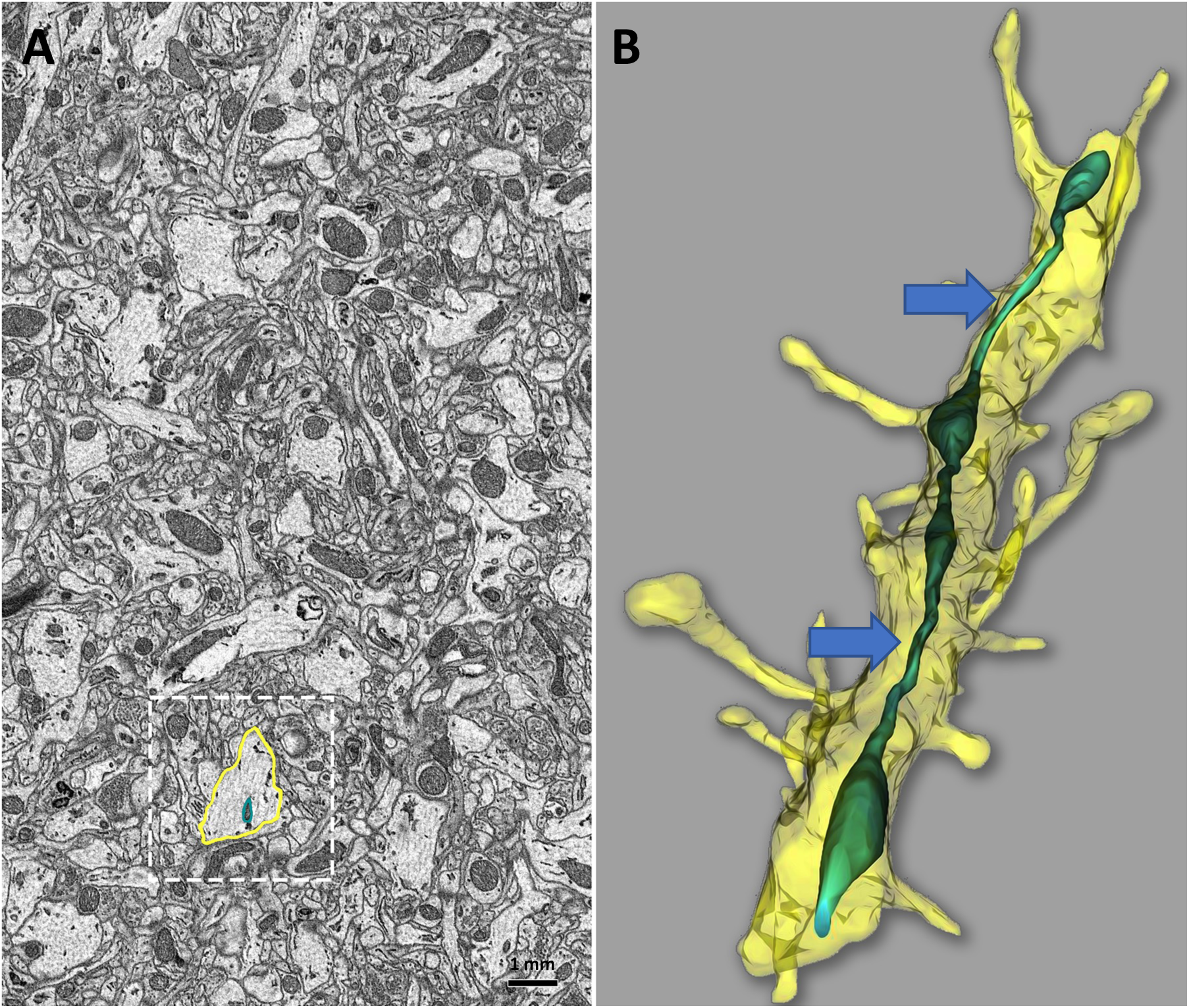
Serial sectioning and 3D reconstruction provide additional insight into ultrastructural details of mouse brain tissue that cannot be fully observed in 2D. Minimal details of a single neuron and mitochondria are observed in a 2D micrograph of a mouse hippocampal tissue of a mouse model of Alzheimer’s Disease, from non-transgenic (NTG1) littermates of a APP/PS1 mouse **A**, boxed area). 3D reconstruction from serial block-face imaging of this same neuron provides far greater detail of neuronal and mitochondrial ultrastructural complexity in boxed area. **B**. Reconstruct utilized for 3D reconstruction allows for better viewing of ultra-structures such as nanotunnels. Scale bar, 1 mm. Blue arrows show nanotunnels.

Further innovations in microscope design are also being developed, which may potentially enhance future data collection techniques. Advances in FIB- and SBF-SEM have increased the accessibility of 3D mitochondrial architecture in the brain and other tissues. The subsequent section summarizes the results of ultrastructural analyses using FIB- and SBF-SEM images obtained from samples prepared using our optimized protocols (see online methods for details), in which expression of OPA1 was reduced.

### SBF-SEM Reveals Increased number of MERC contacts in *Opa1-like* KD *Drosophila* Indirect Flight Muscle and *Opa1* KO Primary Myotubes and Decreased number of MERCs *DRP1-like* KD *Drosophila* Indirect Flight Muscle and *Opa1* KO Primary Myotubes

TEM captures structural MERC changes in 2D. We used SBF-SEM to assess MERC changes in a 3D context, which also allows for a better understanding of cristae dynamics, including changes in volume, shape, and surface area^46^. The apposition of the mitochondrial network with the ER depends on cellular metabolic transitions; therefore, we quantified both loss of *OPA1-like and DRP1-like* in *Drosophila* indirect Flight muscle (Figure 6A–F, SV1–SV2). This revealed significant increases for both measures in the *Opa1-like* knockdown (KD) muscle relative to the WT muscle (Figure 6A–F, Supplemental Video 1–2). Additionally, high-resolution 3D reconstructions revealed increased MERC volumes and lengths in the *Opa1-like*-deficient fly muscles relative to WT muscles while MERC volumes and lengths decreased for *DRP1-like*-deficient fly muscles (Figure 6 G-H). Similarly, these were also measured for myotubes (Figure 6I-N). Relative to control myotubes, MERC length and volume increases were observed in 3D reconstructions of *OPA1*-deficient myotubes while MERC length and volume decreased in *DRP1*-deficient myotubes (Figure 6O–P, Supplemental Video 3–4). Together, these data demonstrated that *OPA1* loss from mouse primary myotubes and *Opa1-like* loss from *Drosophila* muscles increased MERC volume and length and increased MERC tethering. In contrast, DRP-1 loss revealed the opposite effect characterized by decreased MERC volume and length, suggesting a decrease in MERC tethering.

**Figure 6.**
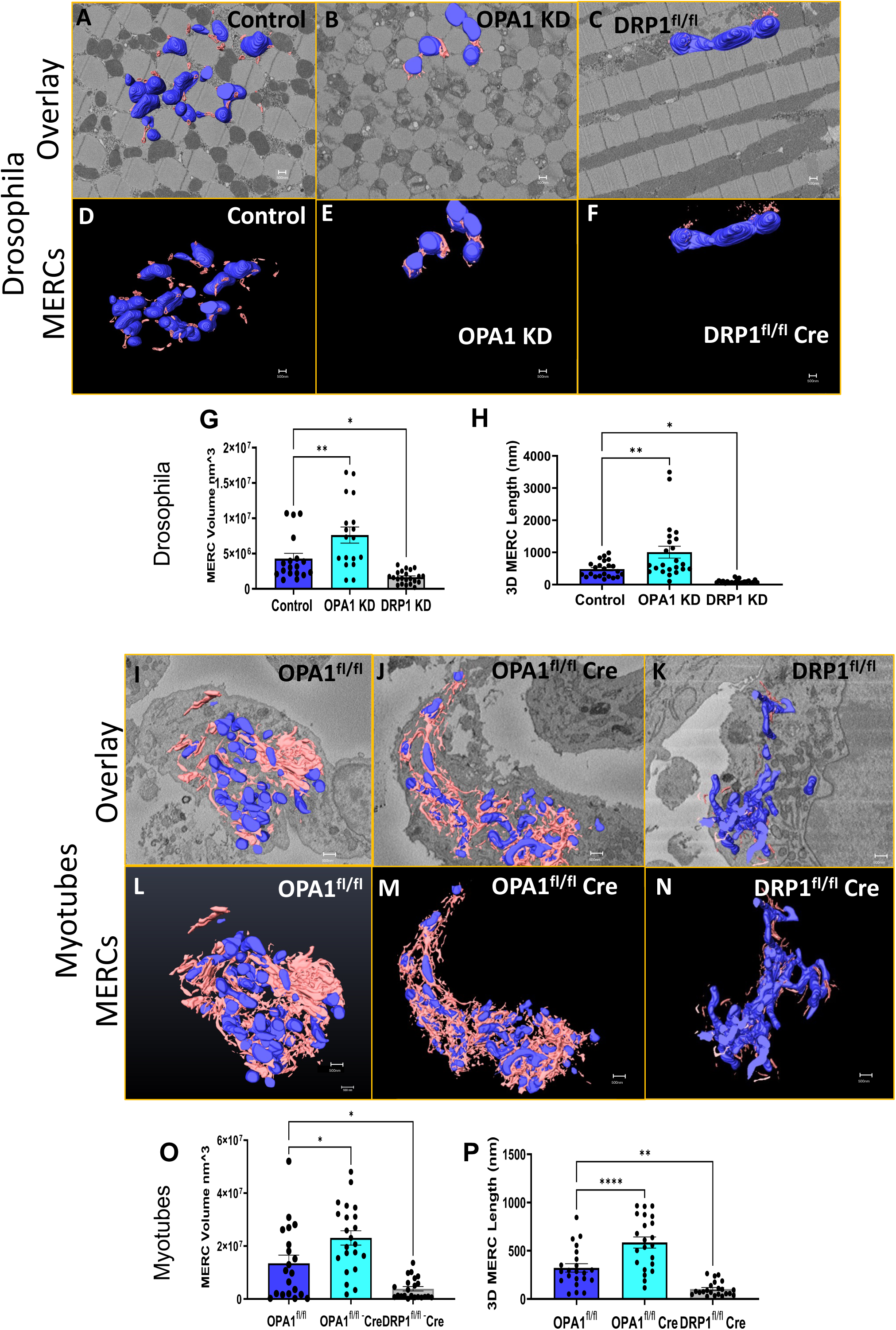
*Opa1-like*-deficiency in skeletal muscle leads to the formation of Mitochondria– Endoplasmic Reticulum (ER) contacts (MERCs) in *Drosophila* and myotubes and *DRP1-like* deficiency reduces MERCs. The 3-dimensional (3D) distribution of single continuous and stationary mitochondria (blue) and endoplasmic reticulum (ER; pink), reconstructed from serial block-face-scanning electron microscopy (SBF-SEM) image stacks of control, *Opa1-like* KD, and *DRP1-like* KD *Drosophila* indirect flight muscle (IFM) fibers **A-F.** and primary myotubes **I-N. G.** Mitochondria–ER contact (MERC) volume and **H.** 3D MERC length, significantly increased in *Opa1-like* KD compared with wild-type (WT) in *Drosophila* skeletal muscle, but all of these metrics decreased in *DRP1-like* KD when compared with the WT. (SV1 [WT]; SV2 [*Opa1-like* KD]). **O.** MERC volume and **P.** 3D MERC length significantly increased in *Opa1-like* KO compared with WT primary myotubes, but decreased in *DRP1-like* KO when compared with the WT (SV3 [WT]; SV4 [*Opa1* KD]). Significance was determined using a non-parametric Welch’s t-test. *P < 0.05, **P < 0.01, ***P < 0.001. SBF-SEM reconstructions from 7 to 23 fully constructed mitochondria, ER, or MERCs. OPA-1 fl/fl represents a control while fl/fl-cre represents deletion of floxed OPA-1.

### Evaluation of Mitochondrial Size and Morphology in *Opa1-like*-Deficient *Drosophila* by SBF-SEM

To assess the role of *Opa1-like* in *the Drosophila* model, we generated mitoGFP (control) flies and *Opa1-like*-deficient flies (Figure 7A–B). Past research has demonstrated changes upon ablation of OPA1^8,14,56,57^. We sought to further elucidate its role in mitochondria remodeling through our methods and usage of 3D SBF-SEM reconstruction. We observed increases in the circularity index and the mitochondrial number (Figure 7C and E) and a decrease in the mitochondrial area (Figure 7D) after *Opa1-like* ablation. We also collected data from the skeletal muscle of WT and Opa1-like deficient flies (Figure 7F–G) and obtained Z-stacks for each condition (Figure 7H–I) to generate a 3D SBF-SEM reconstruction. The mitochondrial morphological reconstruction is shown as viewed virtually from either above (Figure 7J–K) or below (Figure 7L–M) the XY plane. In both cases, following the deletion of *Opa1-like*, there was an overall reduction in mitochondrial volume (Figure 7N). We also observed a decrease in the average 3D mitochondrial length in *Opa1-like* KD *Drosophila* compared with control *Drosophila* (Figure 7O).

**Figure 7.**
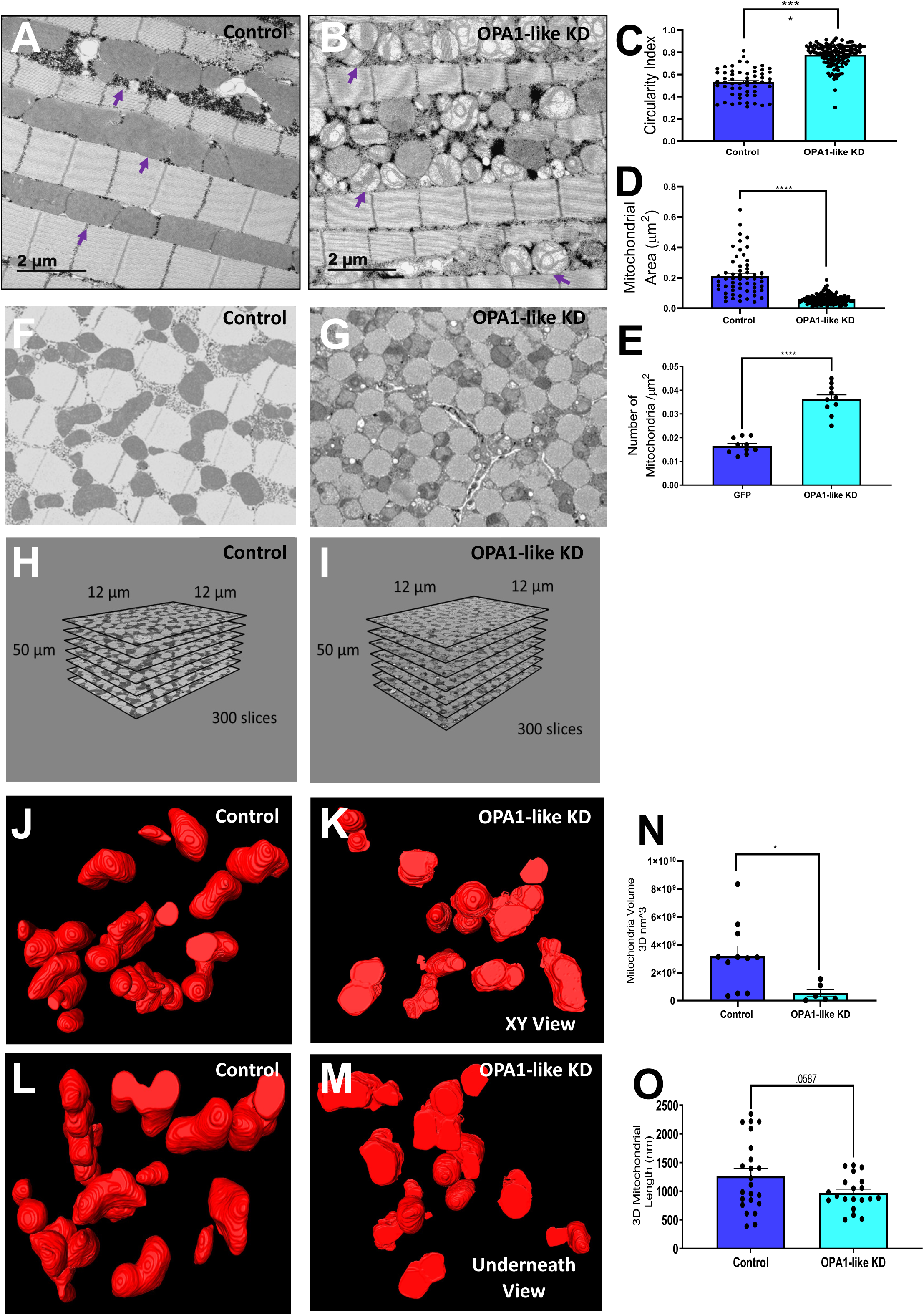
3D reconstruction of control and *Opa1-like* knockdown in *Drosophila*. **A.** Micrograph of mitoGFP (control) and **B,** micrograph of *Opa1-like* knockdown (KD) in *Drosophila* skeletal muscles. Purple arrows identify representative mitochondria. **C.** Circularity index, **D.** mitochondrial area, and **E.** mitochondrial number in control and *Opa1-like*-deficient *Drosophila*. Micrograph of skeletal muscle mitochondrial morphology from **F**., control *Drosophila* and **G**., *Opa1-like* KD *Drosophila.* **H.** Control and **I,** *Opa1-like* KD Z-stack at electron microscopy (EM) resolution from serial block-face-scanning EM (SBF-SEM) used for 3-dimensional (3D) reconstructions. XY 3D reconstruction view of skeletal muscle mitochondrial morphology from **J**., control and **K,** *Opa1-like* KD *Drosophila* using SBF-SEM. Underneath 3D reconstruction view of skeletal muscle mitochondrial morphology from **L,** control and **M,** *Opa1-like* KD *Drosophila* using SBF-SEM. **N.** Average mitochondrial volume and **O.** 3D mitochondrial length in *Opa1-like* KD *Drosophila* skeletal muscle compared with control *Drosophila* using SBF-SEM.

### FIB-SEM of a Gastrocnemius Oxidative Muscle Fiber

A single raw image from an FIB-SEM dataset obtained from a mouse gastrocnemius oxidative muscle fiber is depicted in Figure 8A (100 nm). A 3D rendering of the mitochondrial membranes (red), the ER (green), transverse tubules (t-tubules; blue), lipid droplets (LDs) (cyan), and lysosomes (magenta), partially overlaid on the raw data (gray), is depicted in Figure 8B (100 nm). A raw, magnified image of MERCs, LDs, lysosomes, and t-tubules is shown in Figure 8C (100 nm). Figure 8D shows enhanced magnification images obtained from 3D renderings partially overlaid on the raw data (100 nm). The MERCS (white space) were further visualized in a 3D rendered video (SV 5). Additionally, we illustrated a lone hyperbranched mitochondrion (purple; Figure 8E), a hyperbranched mitochondrion (purple) surrounded by various mitochondria (indicated by an assortment of colors; Figure 8F), and a hyperbranched mitochondrion (green) surrounded by various lysosomes (indicated by an assortment of colors; Figure 8G). These are phenotypes engaged in various organelle-organelle contacts, and specific details of the hyperbranched mitochondria may be important to understand. These volume renderings provide a detailed overview of the organellar structures and their relationships in the oxidative gastrocnemius muscle.

**Figure 8.**
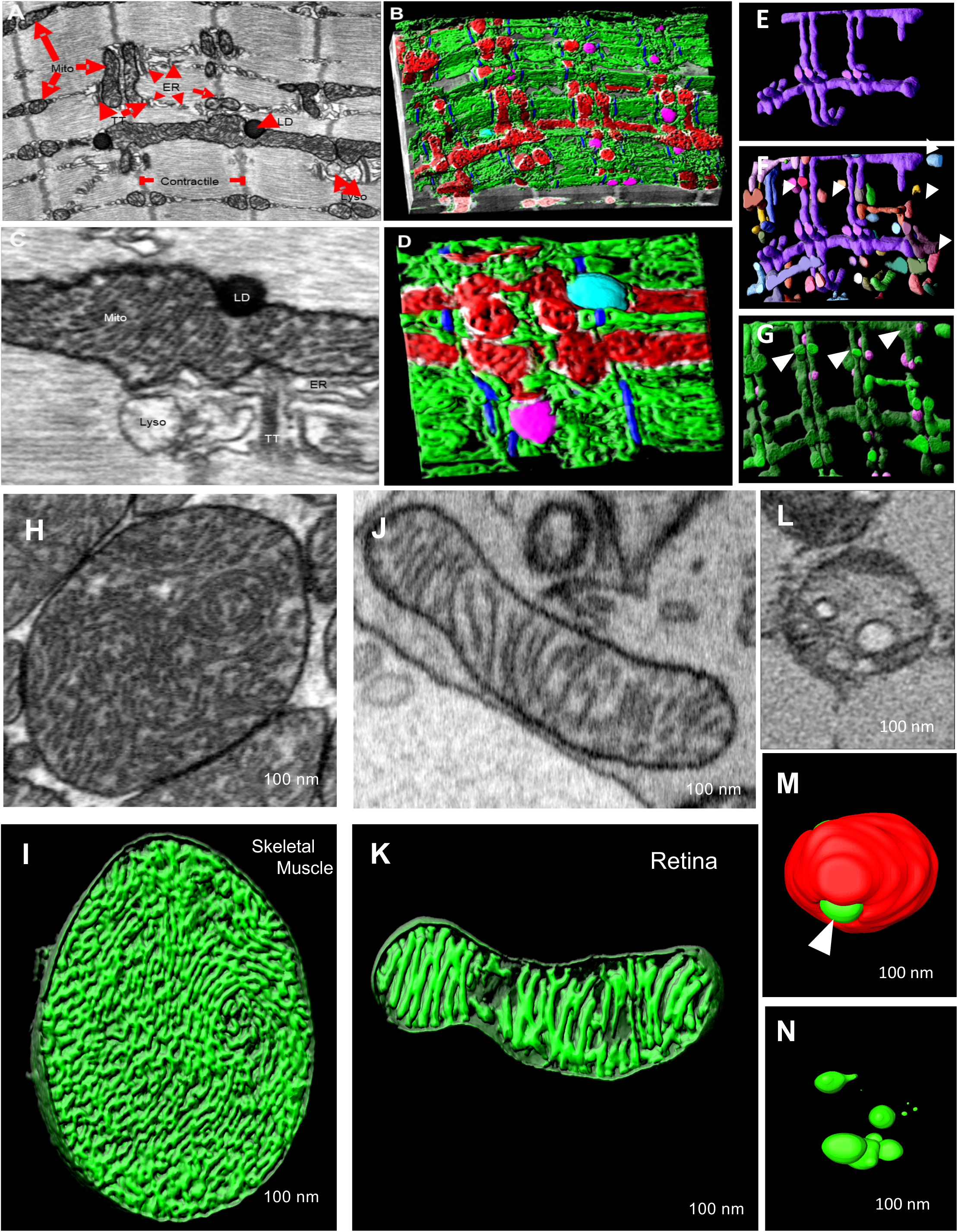
Mitochondrial morphology was visualized using TEM and 3D reconstruction (FIB-SEM and SBF-SEM). **A.** Transmission electron microscopy (TEM) micrograph and **B.** 3-dimensional (3D) reconstruction of mitochondria–endoplasmic reticulum (ER) contacts (MERCs) in mouse skeletal muscle; high magnification (focused ion beam-scanning electron microscopy [FIB-SEM]) **C.** TEM micrograph and **D.** 3D reconstruction of mitochondria (red), actin (blue), lysosome (pink), lipid droplet (teal), MERCs region (white) and ER (green). **E.** 3D reconstruction of a lone hyperbranched mitochondrion from adult mouse skeletal muscle (FIB-SEM). **F.** Hyperbranched mitochondrion surrounded by nearby mitochondria (various colors, white arrowheads) from adult mouse skeletal muscle (FIB-SEM). **G.** Hyperbranched mitochondrion (green) surrounded by lysosomes (pink) from adult mouse skeletal muscle. **H.** Micrograph of cristae-containing mitochondria from WT mouse skeletal muscle. **I.** 3D reconstruction of cristae morphology in WT mouse skeletal muscle using FIB-SEM. **J.** Micrograph of cristae-containing mitochondria from WT mouse retina tissue using FIB-SEM. **K.** 3D reconstruction of cristae morphology in WT mouse retina tissue using FIB-SEM. **L.** Micrograph image of a skeletal muscle mitochondrion in an *Opa1-like* knockdown (KD) in *Drosophila* using SBF-SEM. **M.** 3D reconstruction of cristae (green) with mitochondria (red) in *Opa1-like* KD *Drosophila* skeletal muscle using SBF-SEM. **N.** 3D reconstruction of cristae morphology (green) in *Opa1-like* KD *Drosophila* skeletal muscle using SBF-SEM.

### Resolution of Cristae Morphology by FIB-SEM vs. SBF-SEM

In cases where mitochondria are tethered by an inter-mitochondrial junction, coordinated changes in mitochondrial cristae morphology between distinct organelles can be observed by TEM. Although not all structural cristae details can be observed in 2D images, fine details can be precisely observed using 3D techniques. We examined the efficacy of both 3D imaging techniques, FIB- and SFB-SEM. Specifically, for the resolution of cristae morphology in fine detail and used FIB-SEM technology to progressively explore the sample characteristics.

A single raw image from WT murine skeletal muscle and a 3D rendering of the cristae morphology (green) are illustrated in Figure 8H and I, respectively. Figure 8J and K show the raw images and FIB-SEM 3D reconstructions, respectively, of lamellar cristae morphology from WT murine retinal tissue. We examined the cristae morphology of *Opa1-like* KD *Drosophila* skeletal muscle in both raw and 3D-reconstructed images using SBF-SEM (Figure 8L–N). The XY resolution of SBF-SEM was sufficient to discern cristae resolution (Figure 8M, white arrowhead). In contrast to FIB-SEM, SBF-SEM does not allow for the observation of finer details, such as the lamellar cristae, due to the more limited Z resolution. We also generated a 3D SBF-SEM reconstruction of cristae surrounded by mitochondria in *Drosophila* skeletal muscle (Figure 8N). These data reveal that SBF-SEM could be valuable for assessing the relative organellar relationships in large volumes. FIB-SEM can be used to obtain high-resolution detail in smaller sample volumes.

## Discussion

To realize the full potential of new techniques to assay mitochondrial structures in tissues, sample preparation methods must be reliable and free of artifacts. We used serial sectioning TEM reconstruction methods to develop a protocol that could be used to study mitochondrial morphology in various tissues, including the brain and muscle. MOAS, which forms in response to energetic stress and hypoxia, can easily be missed using conventional 2D EM or fluorescence microscopy techniques^4^. We also demonstrated that procedures that induce hypoxic conditions, including the use of certain anesthetics, perfusion methods, or delayed fixation, are likely to introduce mitochondrial artifacts, such as MOAS. Moreover, our study demonstrated that mitochondrial morphology details in tissues could be more accurately observed using serial sectioning TEM reconstruction (Figure 5) or 3D EM reconstruction (Figure 6-8). We developed a comprehensive protocol for designing experiments to promote the most efficient use of animals and brain tissues. It is suitable for use with both conventional EM techniques based on thick sectioning and advanced techniques based on thin sectioning. In both cases, high-quality micrographs provide rich information, not only in terms of mitochondrial structure but also in terms of other aspects of the cellular environment, including the nuclear shape, the structure of the nuclear pores, the mitochondria–ER network, the sizes and structures of synaptic boutons, and mitochondrial localization within neurites.

The complexity of the intracellular environment is not limited to the mitochondria and can be examined in various tissues, including the brain. This protocol was tested using multiple sectioning techniques, making it suitable for a wide variety of 3D EM reconstruction applications. Depending on the experimental design and the available sample group, the best tissue preparation techniques for EM examination in mice could be achieved through anesthetization by the inhalation of 5% isoflurane/oxygen, followed by cervical dislocation (Figure 4). The use of transcardial PBS perfusion before PFA delays fixation and can result in artifactual changes to the mitochondrial morphology. During direct tissue fixation protocols, using smaller pieces of tissue rather than larger pieces can improve fixation. Adhering to these technical guidelines during the preparation of tissues intended for ultrastructural analyses will avoid the introduction of artifacts and strengthen observations regarding mitochondrial changes. These methods offer reliable steps for the preparation of cells and tissues prior to the analysis of mitochondria and associated structures, including MERCs. MERCs behave as signaling hubs for many molecules, including calcium. Elucidating these interactions may inform the mechanisms underlying diverse pathologies, including neurodegenerative diseases, cancer, diabetes, kidney and liver disease, and developmental disorders^58^. These technical developments could be important for understanding basic cell biology given for example that cristae are crucial for mitochondrial regulation and function, including oxidative phosphorylation^59^. In addition, these techniques can be applied to determine the best practices for imaging other tissues, such as skeletal muscle, and other model organisms, such as flies, cells, and mice (Figures 1–8).

### Measurements of Mitochondrial Cristae

Multiple methods exist for analyzing cristae. The cristae score—a scoring system for cristae quantity and form that ranges from 0 (worst) to 4 (best)—may be an appropriate measurement for use in tissue. The cristae score is effective as it allows for an understanding of whether the cristae structure is intact or has been degraded^10^. However, the cristae score is a subjective judgment that does not incorporate the number of cristae^10^. Thus, the most reliable measurements are cristae volume density, cristae surface area, and cristae number, which are direct measures of changes in the cristae folds or cristae membranes after gene deletion or treatment^5,9^. Mitochondria in tissues can be scored in a similar fashion, using mitochondrial volume density and cristae surface area. These findings can be scored using 2D TEM by examining changes in the typical morphology^5,9,10^. Furthermore, distinguishing which types of cristae are being altered by genes or treatments is also necessary. Cristae may appear as lamellar or tubular. Tubular cristae present a higher base surface area to volume ratio, whereas lamellar cristae are more capable of expansion and have a higher oxidative phosphorylation potential^60,61^. FIB-SEM allows for the 3D reconstruction of the cristae folds, which can be used to distinguish between lamellar and tubular cristae with greater resolution than can be obtained using 2D TEM^5,9,10^.

### Measurement of MERCs

*MFN2* deletion has been reported to increase MERC distance and decrease the ER–mitochondria contact coefficient (ERMICC)^6^, which were confirmed using our quantification methods. We also used the percentage of ER and mitochondria coverage to measure the MERC space (Figure 3)^19^. We found that *MFN2* and *DRP-1* ablation increased the ER–mitochondria distance, and for *MFN2* the ERMICC value indicated a greater percentage coverage of MERCs^6^, consistent with previously published studies^19^ that showed an increased MERC distance.

Mitochondria adjust to metabolic conditions through the parallel remodeling of cristae and MERCs via a mechanism that degrades OPA1. This occurs in a MFN2-dependent manner, suggesting that MERC distance can be altered by changes in energy status^20^. Similar to our findings, some past research showed that *MFN2* ablation increased MERC distance^6,17^. Other studies have reported that the loss of *MFN2* increases MERC coverage using various measurements involving the normalization of the MERC length against either the mitochondrial or ER surface area^19^. Although other studies have suggested a role of DRP1 on MERCs^54^, research showing changes in MERC dynamics upon DRP1 loss is limited. Here, we present a standardized and systematic method for measuring MERCs and quantifying MERC distance, ERMICC, and percent coverage, in addition to performing 3D reconstructions. This protocol can be used to further elucidate MERC dynamics and functions. MERC apposition must be precisely maintained to ensure normal inter-organellar Ca^2+^ transport. Insufficient MERC distance can result in steric hindrance between the components of the Ca^2+^ transporter machinery. Ca^2+^ uptake between the ER and mitochondria is more likely to occur when the organelles are in close proximity, with an ideal distance ranging from 15–30 nm . For example, smaller MERC coverage due to larger MERC distances suggests that changes in response to metabolic conditions would be negligible^1,21^. However, increased MERC coverage associated with increased MERC distance may suggest greater perturbations in the Ca^2+^ uptake or mobilization (the Ca^2+^ transfer effect)^11^. Additionally, smaller MERC distances can allow for the occurrence of lipid transfer^11^. Understanding and accurately quantifying available specific MERC measurements will contribute to understanding changes induced by various treatments. Loss of *MFN2* or *DRP-1* can increase ER stress due to an increase in eukaryotic translation initiation factor 2-alpha kinase (eIF2AK) and protein kinase R (PKR)-like ER kinase (PERK) activity, which are involved in global protein translation attenuation and chaperone expression^23^. These signaling changes can result in ER ballooning.

### Limitations and Considerations

In this study, we demonstrated quantification methods that can be applied to the systematic analysis of organelles in tissues and cells (Figures 2, 3, 5, 6, and S1). While 2D imaging offers important information, it does not capture the full mitochondrial dynamics that 3D reconstruction captures. Therefore, in addition to 2D imaging, we suggest using FIB-SEM^21,24,28,33,37,41,42,62,63^ or confocal analysis^5^ to determine mitochondrial volume. To measure the entire cell, we recommend 3D reconstruction, which describes the whole cell and a reasonable overview of the mitochondrial number. Unless dealing with a limited data set, we suggest using a histogram for mitochondrial area. This allows for examination of shifts in the distribution of the mitochondrial area and better understanding the heterogeneity across mitochondria areas^5^. Additionally, a histogram can provide insights regarding mitochondrial size, which may be altered in response to gene deletion or treatment. Other mitochondrial calculations may not be as viable, which was observed during the 3D reconstruction. For example, when assessing the number of mitochondria per cell or the total mitochondrial volume, we found that the limited number of mitochondria included in each plane can be a limitation. However, estimating volume may still be performed without the resolution needed to to count individual mtiochondria. In general, the total mitochondria may be a limiting quantification, and 3D reconstruction can limit the number of mitochondria quantified due to the associated time it takes.

Other tools, in addition to the imaging techniques detailed above, can be used to evaluate MERC tethering, such as the proximity ligation assay (PLA). PLA permits the detection of protein-protein interactions in situ at distances of less than 40nm at endogenous protein levels. Co-immunoprecipitation (Co-IP), ER Tracker, and MitoTracker analyses can also be used to examine changes in MERC colocalization in the absence or presence of genes of interest and treatment conditions^17,19,22,43^. ER Tracker is an ER-specific dye that shares some overlap with mitochondria. Mitochondrial staining can be used to confirm differences in ER/mitochondrial colocalization with changes in TEM analyses. MERC distance can be measured by examining the ER–mitochondria contact space distance or coverage using FIB-SEM^1^, or electron tomography^21^. Immunogold labeling^22^ can also be used by individually staining proteins known to be associated with MERC tethering or mitochondria. The colocalization of these immunogold-labeled dots can also be examined. This technique can also be harnessed to validate changes in MERC tethering proteins content^21^, analogous to insights gained from PLA^43,55^.

## Conclusion

The present study demonstrates an optimized approach for preserving specimens and measuring organelle morphology using 2D and 3D imaging. We demonstrate that methods that create hypoxic conditions could introduce unintentional artifacts and describe optimal conditions for fixing cells in culture to preserve cellular and mitochondrial integrity. Furthermore, this study presents standardized quantification methods that can be used to measure mitochondrial morphology. It can further be used to measure other organellar structural features, including mitochondrial size and MERCs. Finally, we verified previously described phenotypes to illustrate the efficacy of our methodology. Using these methods, investigators should be able to accurately and reproducibly visualize and measure ultrastructural changes in cells and tissues using TEM imaging techniques.

## Limitations of the study

Although our study covers cell lines, *Drosophila melangaster* and *Mus musculus*, this has not been looked at in other organisms, such as *Caenorhabditis elegans* and *Danio rerio*, and is unclear if these techniques are consistent. Additionally, images for showing artifacts have been shown through TEM (2D) and does not show morphological changes in 3D.

## Supplementary Figures

**Figure S1 Quantification of mitochondrial characteristics in *OPA1*-deficient skeletal muscle myoblasts and *MFN2*-deficient or *OPA1*-deficient skeletal muscle myotubes**

**A.** Western blotting, with GAPDH as a control, showed successful knockout of DRP-1, MFN-2, and OPA-1. Qualitative polymerase chain reaction showed a fold decrease of more than 50% in the case of **B.** OPA-1 **C.** DRP-1 and **D.** MFN-2. Ultrastructure of (**E-G),** control and (**H-J),** *DRP-1*-deficient primary skeletal muscle myotubes. **K.** Mitochondria length, **L.** mitochondria area, **M.** Cristae number, **N.** Cristae number, and **O.** Cristae surface area in control and *DRP-1*-deficient primary skeletal muscle myotubes.

**Figure S2 Mitochondria-Endoplasmic Reticulum Contact Site (MERC) ultrastructure in *DRP-1*-deficient primary skeletal muscle myotubes.**

Ultrastructure of (**A-B),** control and (**C-D),** *DRP-1*-deficient primary skeletal muscle myotubes. There was a large increase in **E.** DRP1-KO area of MERCs, and **F.** an increase in MERC distance in DRP-1 KO.

**Supplementary Video 1.** Representation of the 3D reconstruction of MERCS in wild-type *Drosophila* muscle.

**Supplementary Video 2.** Representation of the 3D reconstruction of MERCS in *OPA1* knockdown *Drosophila* muscle.

**Supplementary Video 3.** Representation of the 3D reconstruction of MERCS in *OPA1^fl/fl^* myotubes.

**Supplementary Video 4.** Representation of the 3D reconstruction of MERCS in *OPA1^fl/fl^-Cre* myotubes.

**Supplementary Video 5.** Representation of the 3D reconstruction of mitochondria membranes (red), ER (green), actin (blue), LDs (cyan), lysosomes (magenta), and MERCs (white space).

## STAR Methods (provides all technical details necessary for the independent reproduction of the methodology)

### Animals

#### Mice

All procedures were performed using humane and ethical protocols approved by the Institutional Animal Care and Use Committees of the Mayo Clinic and the University of Iowa, in accordance with the National Institute of Health’s Guide for the Care and Use of Laboratory Animals. Mice were housed on a 12-hour light/dark cycle and were provided access to food and water. C57BL/6 (wild-type; WT) female mice of various ages were used in the study, with four to six animals examined for each condition. For definitive MOAS identification (positive control), APPSWE (human APP 695 gene containing the double mutations: K670N, M671L) female mice (n = 3), a model of familial AD, were used (Hsiao, 1996). A total of 16 male mice were used to isolate primary satellite cells from control littermates (*lox-OPA-1* mice), and another 8 mice each were used as control littermates (*lox-OPA-1* mice) and *OPA-1-HSA-CreER^T^*^2^. All mice were on a pure C57BL/6J genetic background.

### Drosophila Strains and Genetics

Genetic crosses were performed on a yeast corn medium at 22 °C unless otherwise stated. *Mef2-Gal4* (III) was used to drive the muscle-specific Opa-1-like (OPA1) (BS #32358) knockdown (KD). *Tub-Gal80ts* (BS #7019) and *Mef2 Gal4* (BS #27390) were used for the conditional muscle-specific *Opa-1-like* KD. Genetic crosses were set up at 18 °C and then shifted to 29 °C at the larval stage (L3). *Mef2 Gal4; UAS-mito-GFP* (II chromosome) (BS #8442) was used as control. Stocks were obtained from the Bloomington Drosophila stock center. All chromosomes and gene symbols are as described in FlyBase (http://flybase.org).

### Primary Cell Culture

Satellite cell isolation was performed as previously described (Pereira et al., 2017)^47^. Satellite cells from *OPA-1^fl/fl^* mice were plated on BD Matrigel-coated dishes and activated to differentiate into myoblasts in Dulbecco’s modified Eagle medium (DMEM)-F12 containing 20% fetal bovine serum (FBS), 40 ng/ml basic fibroblast growth factor, 1× non-essential amino acids, 0.14 mM β-mercaptoethanol, 1× penicillin/streptomycin, and Fungizone. Myoblasts were maintained with 10 ng/ml basic fibroblast growth factor and differentiated in DMEM-F12 containing 2% FBS and 1× insulin–transferrin–selenium when 90% confluency was reached. Three days after differentiation, myotubes were infected with an adenovirus expressing GFP-Cre to achieve OPA-1 deletion. Adenoviruses were obtained from the University of Iowa Viral Vector Core facility. Experiments were performed between 3 and 7 days after infection.

### Measuring Organelle Morphology

To perform unbiased morphometric analysis of organelles, separate individuals should be responsible for conducting the fixation and image acquisition, while a group of blinded individuals should be responsible for quantification^44^. The use of ImageJ, an open-source image processing software developed by the National Institutes of Health (NIH) and designed to analyze multidimensional scientific images, such as TEM and confocal microscopy data sets, is suitable for the quantification analysis of metrics, including length, area, perimeter, and circularity index. Circularity index specifically is a measure of roundness calculated by 4 π * (A/P^2) where a is the area and p is the perimeter of the desired organelle^44,64^. The entire cell is divided into quadrants, using the ImageJ plugin *quadrant picking* (https://imagej.nih.gov/ij/plugins/quadrant-picking/index.html), and all measurements should be performed with a minimum of 10 cells (n = 10) in each of three independent analyses^44^. For these specific quantification methods, please refer to Lam et al. ^44^.

## Technical Methods

### TEM Processing of Skeletal Muscle Myoblasts

To delineate essential calculations for analyzing TEM images, we investigated mitochondrial and other organelle morphology in human, primary mouse, and immortalized mouse skeletal myoblasts. Skeletal muscle satellite cells were isolated, cultured to the myoblast state, and placed in six-well poly-D-lysine–coated plates for TEM processing. Media was substituted with 2.5% glutaraldehyde in 0.1 M sodium cacodylate buffer (pH 7.2) at 37 °C and incubated for 5 minutes, 10 minutes, 60 minutes, or 24 hours depending on the trial. After our findings in Figure 1, all subsequent trials employed 24 hours. The remainder of the process was performed directly in the culture plate at room temperature. After fixation, cells were rinsed twice with 0.1 M sodium cacodylate buffer (pH 7.2) for 5 minutes each. Next, a secondary fixation was performed using 1% osmium tetroxide + 1.5% potassium ferrocyanide in 0.1 M sodium cacodylate buffer (pH 7.2) for 30 minutes. The plate was washed repeatedly with 0.1 M sodium cacodylate buffer (pH 7.2) until the liquid appeared colorless. The plate was then washed twice with the same buffer and twice with deionized water (diH_2_O) for 5 minutes each. Subsequently, the plate was incubated in an en-bloc staining solution of 2.5% uranyl acetate overnight. Dehydration was performed the following day using a graded ethanol series—25%, 50%, 75%, 95%, and two changes of 100% ethanol for 5 minutes each. Infiltration with an epoxy resin, Eponate 12^™^ (Ted Pella Inc, cat # 18005), was performed by gently and thoroughly mixing Eponate 12™ with 100% ethanol in a 1:1 solution, replacing the dehydrant with the mixture and incubating for 30 minutes. The infiltration procedure was repeated three times using 100% Eponate 12^™^ for at least 1 hour each. Finally, the intact plate was placed in fresh media in a 70 °C oven for at least 24 hours to cure.

After hardening, the plates were removed from the oven, and the resin-embedded cells were separated from the plastic by cracking the plate and immersing the plate in a liquid nitrogen bath. The temperature differential caused by removal from the bath created a gas layer between the cell-embedded resin and the plate, which was then broken. A jeweler’s saw was used to cut an en-face block fitting into the Leica UC6 ultramicrotome (Leica Biosystems) sample holder. The section was then placed onto formvar-coated (Electron Microscopy Sciences, cat # 15810) copper grids. The grids were counterstained in uranyl acetate and Reynold’s lead citrate for 2 minutes each. The samples were imaged using a JEOL 1230 transmission electron microscope (JEOL Ltd.) at 120,000× with an accelerating voltage. This protocol can also be used to process other cell types, such as hepatocytes, endothelial cells, adipocytes, and cardiomyoblasts.

### TEM Processing of Skeletal Muscle Myotubes

Differentiation of myoblasts on poly-D-lysine–coated plates alters the normal morphology of skeletal muscle myotubes. Therefore, we employed Matrigel coating, which allowed for the even distribution and proper morphology of skeletal muscle myotubes. To obtain skeletal muscle myotubes, skeletal muscle satellite cells were isolated, cultured to the myoblast state, and split on Matrigel-coated plates for myotube formation. The processing of cells grown in a Matrigel-coated plate requires changes to the previously described protocol.

First, 2.5% glutaraldehyde in 0.1 M sodium cacodylate buffer (pH 7.2) was warmed to the temperature of the cell culture media. Cell media was replaced with the warmed fixative and timed for 5 minutes, 10 minutes, 60 minutes, or 24 hours depending on the trial. After our findings in Figure 1, all subsequent trials employed 24 hours. All the following steps were performed directly in the culture plate at room temperature, with gentle agitation on a rotator during incubations to minimize the disturbance of the cell layer. After fixation, the cells were rinsed at least three times with 0.1 M sodium cacodylate buffer (pH 7.2) for 10 minutes each. Secondary fixation was performed using 1% osmium tetroxide in 0.1 M sodium cacodylate buffer (pH 7.2) with 1.5% potassium ferrocyanide for 45 minutes. Washing was performed with 0.1 M sodium cacodylate buffer (pH 7.2) until the liquid appeared colorless, followed by two washes with the same buffer and three washes with diH_2_O for 5 minutes each. En-bloc staining using 2.5% uranyl acetate was performed overnight. Dehydration was performed the following day with a graded series of ethanol at 25%, 50%, 75%, 95%, and two changes of 100% ethanol for 10 minutes each while ensuring that the cells did not dry out. Infiltration was performed with an epoxy resin, Eponate 12™ (Ted Pella Inc, cat # 18005) by gently and thoroughly mixing Eponate 12™ with 100% ethanol in a 1:1 solution and replacing the dehydrant with the mixture, allowing infiltration for 1 hour. Cells were additionally infiltrated with three incubations in 100% Eponate 12™ for at least 2 hours for each change of solution. One of the changes was performed overnight at 4 °C. Finally, the intact plate was placed in fresh media in a 70 °C oven for at least 24 hours to cure. Extraction from the plates, cutting, staining, and microscopy were performed as described in the preceding section.

### Focused Ion Beam-Scanning Electron Microscopy (FIB-SEM) Processing of Adult Skeletal Muscle Fibers

To reveal muscle cell organelle connectivity in 3D, we analyzed the morphology and interactions between mitochondria, ER, LDs, lysosomes, and t-tubules in mouse gastrocnemius muscle using a protocol that highlights the organelle membranes within the cell. Male C57BL/6J mice were placed on a heated bed and anesthetized via inhalation of 2% isoflurane. Hindlimb hair and skin were removed, and hindlimbs were immersed in 2% glutaraldehyde in 100 mM phosphate buffer, pH 7.2. After 30 minutes, the gastrocnemius muscle was removed, cut into 1 mm^3^ cubes, and placed in 2.5% glutaraldehyde, 1% paraformaldehyde, 120 mM sodium cacodylate, pH 7.2–7.4 for 1 hour. After five 3-minute washes with 100 mM cacodylate buffer at room temperature, samples were placed in 3% potassium ferrocyanide, 200 mM cacodylate, 4% aqueous osmium for 1 hour on ice, washed five times for 3 minutes each in diH_2_O, and incubated for 20 minutes in fresh thiocarbohydrazide solution at room temperature. Samples were then incubated on ice for 30 minutes in 2% osmium solution and washed five times for 3 minutes each in bi-distilled H_2_O. The sample was then incubated overnight in 1% uranyl acetate solution at 4 °C, washed five times for 3 minutes each in bi-distilled H_2_O, incubated in 20 mM lead nitrate, 30 mM aspartic acid, pH 5.5 at 60 °C for 20 minutes, and washed five times for 3 minutes each in bi-distilled H_2_O at room temperature. The sample was next incubated sequentially in 20%, 50%, 70%, 90%, 95%, and 100% ethanol for 5 minutes each, incubated in 1:1 Epon:ethanol solution for 4 hours, and incubated in 3:1 Epon:ethanol at room temperature overnight. The next day, samples were incubated sequentially in fresh 100% Epon for 1 hour and 4 hours. After removing excess resin using filter paper, the samples were placed on aluminum Zeiss SEM Mounts (Electron Microscopy Sciences, #75510) in a 60 °C oven for 2 days. Stubs were then mounted on a Leica UCT Ultramicrotome (Leica Microsystems Inc., USA) and faced with a Trimtool 45 diamond knife (DiATOME, Switzerland), with a feed of 100 nm at a rate of 80 mm/s.

FIB-SEM images were acquired by a Zeiss Crossbeam 540 using Zeiss Atlas 5 software (Carl Zeiss Microscopy GmbH, Jena, Germany) and collected using an in-column energy selective backscatter with filtering grid to reject unwanted secondary electrons and backscatter electrons up to a voltage of 1.5 kV at a working distance of 5.01 mm. Milling was performed by an FIB, operating at 30 kV, with a 2–2.5 nA beam current and 10 nm thickness. Image stacks were aligned using Atlas 5 software (Fibics) and exported as TIFF files for analysis. This protocol, designed for evaluating organelle connectivity in adult skeletal muscle fibers, also works for cardiomyocytes, neonatal myocytes, and brain tissues^20^.

### Analysis of FIB-SEM Images for Adult Skeletal Muscle Fibers

To evaluate 3D organelle connectivity within the FIB-SEM volumes, we first separated each type of cellular structure within the greyscale datasets. After normalizing contrast throughout the dataset using the 3D Enhance Local Contrast tool in ImageJ (NIH, Bethesda, MD, ImageJ.net) and saving it as an HDF5 (Hierarchal Data Format 5) file, organelle segmentation was completed with the Pixel Classification module in the Ilastik software package^6^ (Ilastik.org). The raw HDF5 file was imported into the software, and the features used to train the pixel classifier were as follows: Color/Intensity 0.3–10 pixels; Edge 0.7–10 pixels; and Texture 0.7–10 pixels. Training of the pixel classifier was performed by tracing the organelle and contractile structures in the middle XY, XZ, and YZ planes of the volume. Training labels were created for the mitochondrial outer membrane, mitochondrial interior, LDs, sarcoplasmic reticulum, t-tubules, lysosome membrane, lysosome interior, and the sarcomeric I-bands, A-bands, and Z-disks. After tracing, the Live Update tool was used to compare the segmentation results to the raw data to check for errors. When needed, further training iterations were performed to reduce errors. Pixel probability maps were then exported as 8-bit HDF5 files.

Segmentation of individual mitochondria and lysosomes was more accurate than initial bulk segmentation; thus, additional training was performed. This second step was performed using the MultiCut module in Ilastik. Raw data was loaded for pixel classification, and then the HDF5 file containing the outer mitochondrial membrane or lysosome membrane probabilities was loaded. Superpixels were created using the membrane probabilities as the input channel, a threshold of 0.2, a presmooth before seeds value of 1.0, clustered seed labeling, and default other values. The training was then performed by marking mitochondrial or lysosomal boundaries in red and non-mitochondrial or lysosomal boundaries in green and selecting the means of the raw data and membrane probabilities standard (edge) and standard (sp) as features. The Live Predict tool was used to create initial boundary predictions before using the Live Multicut tool with the Nifty_FmGreedy solver and 0.5 beta to generate the initial individual mitochondrial or lysosomal segmentations. Increased accuracy was obtained by additional boundary training and updating the multicut, when necessary. The multicut segmentation was then exported as an 8-bit HDF5 file for evaluation. Notably, this image segmentation routine works well for cardiomyocytes, neonatal myocytes, myoblasts, satellite cells, brain, retina, and red blood cells, among other tissues^20^. 3D renderings of segmented cellular structures were performed using the Volume Viewer plugin in ImageJ.

### Serial Block-Face-Scanning Electron Microscopy (SBF-SEM) Processing of Drosophila Muscle Fibers

Samples for serial block-face scanning electron microscopy (SBFSEM) were prepared using a procedure based on that developed by Deerinck^1^. Briefly, fresh sample was fixed by immersion in 2% glutaraldehyde + 2% paraformaldehyde in 0.1 M cacodylate buffer containing 2 mM calcium chloride. After fixation, sample was rinsed in 0.1 M cacodylate buffer and placed into 2% osmium tetroxide + 1.5% potassium ferracyanide in 0.1 M cacodylate, washed with nH_2_O, incubated at 50°C in 1% thiocarbohydrazide, incubated again in 2% osmium tetroxide in nH_2_O, rinsed in nH_2_O and placed in 2% uranyl acetate O/N. The next day, sample is rinsed again in nH_2_O, incubated with Walton’s lead aspartate, dehydrated through an ethanol series, and embedded in Embed 812 resin. Based on rOTO stains developed by Willingham and Rutherford^2^, this procedure introduces a considerable amount of electron dense heavy metal into the sample to help provide the additional contrast necessary in SBFSEM.

To prepare embedded sample for placement into the SBFSEM, a ∼1.0 mm^3^ piece was trimmed of any excess resin and mounted to an 8 mm aluminum stub using silver epoxy Epo-Tek (EMS, Hatfield, PA). The mounted sample was then carefully trimmed into a smaller ∼0.5 mm^3^ tower using a Diatome diamond trimming tool (EMS, Hatfield, PA) and vacuum sputter-coated with gold palladium to help dissipate charge.

Sectioning and imaging of sample was performed using a VolumeScope 2 SEM^TM^ (Thermo Fisher Scientific, Waltham, MA). Imaging was performed under low vacuum/ water vapor conditions with a starting energy of 3.0 keV and beam current of 0.10 nA. Sections of 50 nm thickness were cut allowing for imaging at 10 nm x 10 nm x 50 nm spatial resolution^65–67^.

Image analysis, including registration, volume rendering, and visualization were performed using *ImageJ*^3^, *Reconstruct*, and *Amira* (Thermo Fisher) software packages.

### Serial Block-Face-Scanning Electron Microscopy (SBF-SEM) Processing of Retina Tissue

Retina tissues were isolated via freeze capture^68^. Liquid propane was chilled with dry ice for approximately 15 minutes before being inverted to ensure a liquid jet. In tandem, 20 mL of solvent composed of 97% methanol and 3% acetic acid was also chilled with dry ice. Once mice were euthanized via the methods outlined above, the eyes were rapidly removed, immersed in chilled propane for 1 min, and quickly transferred to the chilled solvent. From there, sections of tissue 4-µm-thick were obtained using a Leica RM2125Rt microtome.

## Acknowledgements

We would like to thank our undergraduate colleagues Benjamin Kirk and Innes Hicsasmaz for assisting with the optimization of this analysis technique. This work was supported by NIH grants R01HL108379 and R01DK092065 to E.D.A; NIH NIA RF1AG55549, NIH NINDS R01NS107265, RO1AG062135, AG59093, AG072899 to E.T.; T32 HL007121, the UNCF/BMS EE Just Faculty Fund, Career Award at the Scientific Interface (CASI Award) from Burroughs Welcome Fund (BWF). ID # 1021868.01, the Ford Foundation, NIH SRP subaward to #5R25HL106365-12 from the NIH PRIDE Program to A.H. Jr; and NSF MCB #2011577I to S.A.M; 2U54CA 1633069-07 from NCI and R25 GM059994/GM/NIHGMS NIH to H.K.B. Its contents are solely the responsibility of the authors and do not necessarily represent the official view of the NIH. The funders had no role in study design, data collection and analysis, decision to publish, or preparation of the manuscript.

## Author contributions

Conceptualization, A.J.H., P.K., T.A.C., J.L.S., R.O.P., B.G., E.T., and E.D.A. Methodology, A.J.H., P.K., T.A.C., M.M., J.S., L.Z., S.T., A.J., R.E.G., E.G.L., Z.V., C.K.E.B., I.H., R.A.C.E., A.D., J.L.S., B.G., E.T., and E.D.A. Investigation, A.J.H., P.K., T.A.C., M.M., J.S., L.Z., S.T., A.A., A.J., R.E.G., E.G.L., Z.V., J.P., C.K.E.B., I.H., R.A.C.E., A.D., Y.D.K., S.B., J.L.S., B.G., E.T., and E.D.A. Writing – Original Draft, A.J.H., P.K., T.A.C., A.A., K.N., M.B., E.G.L., Z.V., H.K.B., A.G.M., J.P., C.K.E.B., I.H., S.A.M., R.A.C.E., A.D., Y.D.K., J.L.S., B.G., E.T., and E.D.A. Writing – Review & Editing, A.J.H., P.K., T.A.C., K.N., M.B., E.G.L., Z.V., H.K.B., A.G.M., C.K.E.B., S.A.M., R.A.C.E., S.B., J.L.S., B.G., E.T., and E.D.A. Funding Acquisition, A.J.H., P.K., T.A.C., B.G., E.T., and E.D.A. Resources, A.J.H., P.K., T.A.C., J.S., J.L.S., R.O.P., B.G., E.T., and E.D.A. Supervision, A.J.H., P.K., T.A.C., J.S., J.L.S., R.O.P., B.G., E.T., and E.D.A.

