## Supplementary figures and images for "A comprehensive approach to artifact-free sample preparation and the assessment of mitochondrial morphology in tissue and cultured cells"

### Supplemental Figures

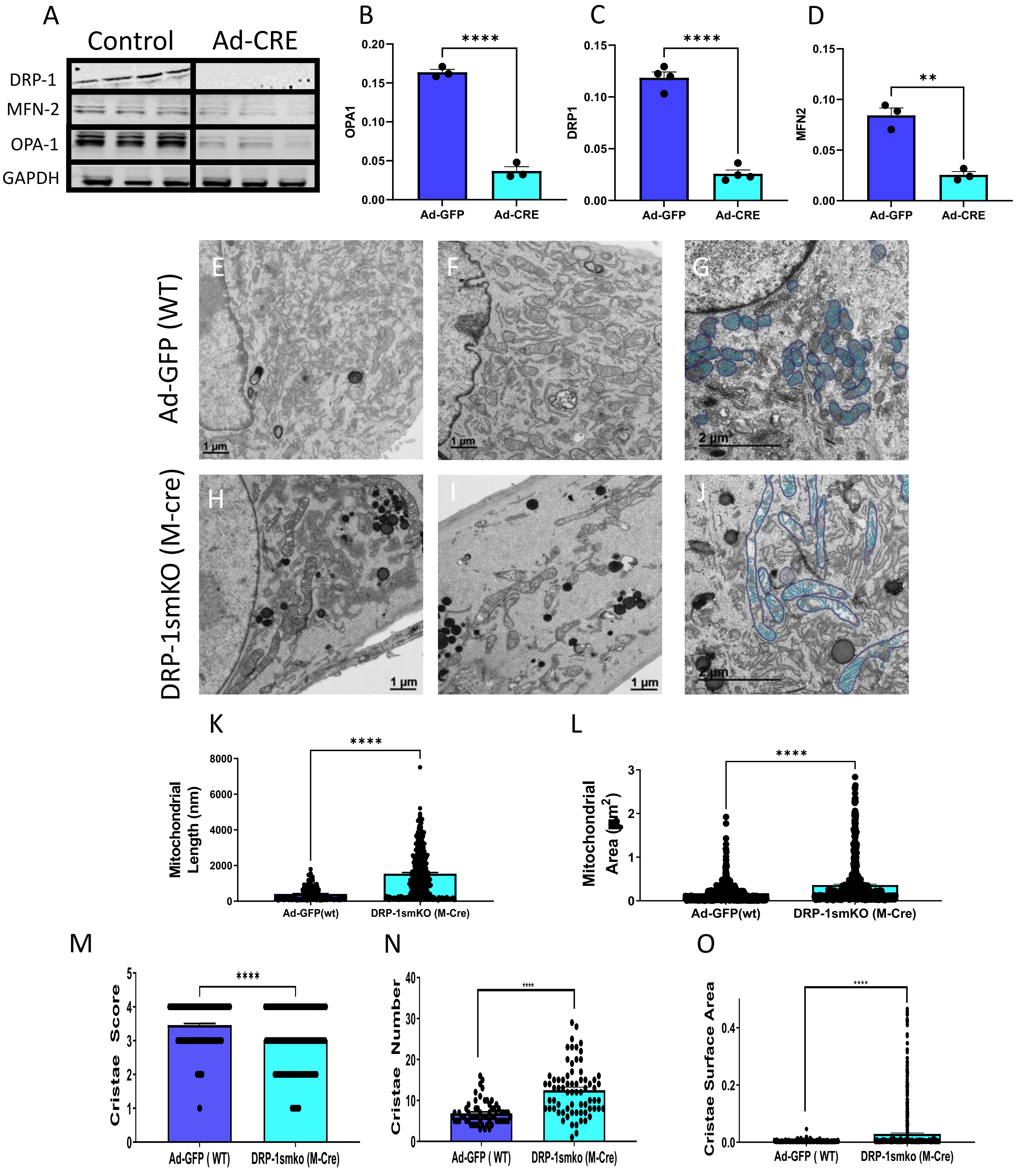

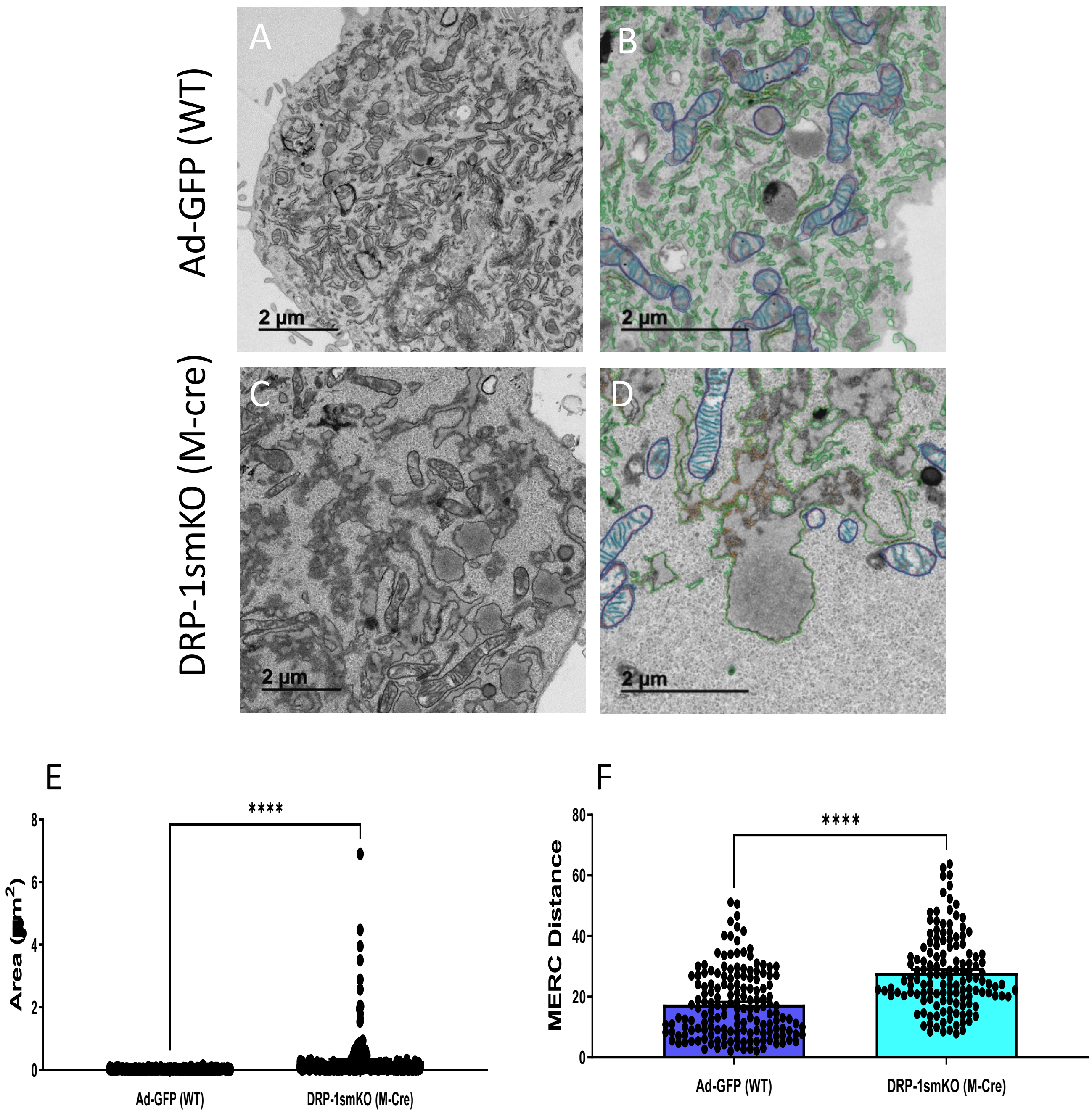
